# The apoptosis gene *BCL-X* splice isoforms have opposing effects in diabetic kidney disease: potential treatment target and prognostic value

**DOI:** 10.1101/2025.06.06.658276

**Authors:** Megan Stevens, Monica L Ayine, Kim Gooding, Angela Shore, Pedro Marqueti, Sebastian Oltean

**Affiliations:** Department of Clinical and Biomedical Sciences, University of Exeter Medical School, Exeter, United Kingdom; NIHR Exeter Clinical Research Facility, Royal Devon University Healthcare Foundation Trust, Exeter United Kingdom

**Keywords:** diabetic nephropathy, alternative splicing, BCL-X, apoptosis, therapeutic targets

## Abstract

In recent years, the importance of alternative splicing (AS) of certain genes in the nature and progression of diabetic nephropathy (DN) has been studied. We report a novel AS event observed in the diabetic kidney – AS of the apoptosis gene *BCL-X* to increase the pro-apoptotic *BCL-XS* and decrease the anti-apoptotic *BCL-XL*. This study aimed to further investigate the role of this novel AS event in the pathogenesis of DN.

To characterize important splicing events in progression of DN, human glomerular endothelial cells (GEnCs) were exposed to a diabetic environment for 1 week and RNAseq was performed. While several splicing changes have been discovered we have focused further on the apoptosis gene *BCL-X* isoforms.

In GEnCs, an upregulation of the *BCL-XS/BCL-XL* ratio was observed which resulted in an increase in GEnC apoptosis. An upregulation of *IL-6* was also observed; treatment with IL-6 alone induced a dose-dependent shift in *BCL-X* splicing to promote expression of the pro-apoptotic *BCL-XS*. Furthermore, we identified certain splicing factors, SF3B1 and PTBP1, involved in the *BCL-X* splicing regulation. A switch in expression of the anti-apoptotic *BCL-XL* isoform rescues apoptosis, suggesting a possible therapeutic avenue. In patients, an increase in the pro-apoptotic *BCL-XS* in urinary RNA correlated with a decline in the GFR, while in blood it correlated with the level of albuminuria.

There is an increase in the pro-apoptotic *BCL-XS* isoform in the diabetic glomerular endothelium, resulting in increased GEnC apoptosis. Increased *BCL-XS* expression correlates with a markers of renal function decline, implicating this AS event as a potential biomarker for DN severity and suggesting that the switch of isoforms from *BCL-XS* to *BCL-XL* may for a novel therapeutic strategy.

## 1. Introduction

With the increasing incidence of diabetes worldwide, there has been an increase in diabetes-associated complications, including diabetic nephropathy (DN). A large proportion of patients with DN will eventually progress to end-stage renal failure and require long-term kidney replacement treatment (dialysis). Due to the large number of such patients, there is a considerable, and increasing, strain on health system resources worldwide. Therefore, new treatments that can slow the progression of DN are urgently needed. (1–3).

At present, clinicians have a few therapeutic tools to use for this purpose. Blood pressure lowering angiotensin-converting enzyme (ACE) inhibitors have long been proven to be beneficial in decreasing proteinuria and slowing the degradation of kidney function. Furthermore, more recently, SGLT2 inhibitors designed to lower glycaemia have shown remarkable effects in the protection of kidney protection function; while the mechanism is not yet completely understood, it is clearly not due only to decreasing glycaemia. However, the above-mentioned drugs, although extremely useful, were not purposely designed for the treatment of DN; their effects were discovered serendipitously. It is therefore thought that understanding the molecular mechanisms involved in the progression of DN and designing drugs to directly address these mechanisms has the potential to elicit new classes of drugs with good efficiency in slowing down the progression of DN (4).

One new area of therapeutic research in various diseases is the modulation of alternative splicing (AS). AS has been shown to occur in over 94% of genes in humans and therefore represents an important level in deciding the proteome diversity of the functional capabilities of cells (5, 6). Since AS is so widespread, it is not surprising that virtually all diseases display aberrant splicing, which includes the appearance of new, specific disease isoforms or modifications to the ratios of existing isoforms (7, 8). Either way, AS has the potential to maintain and drive the pathogenic process. Therefore, a new research field is looking into manipulation of these isoforms for therapeutic benefit, for example, by switching ratios of splice isoforms to their normal rapport (9).

To explore these therapeutic AS decisions further, we need to understand the potentially pathogenic isoforms. Therefore, this study aimed to determine important AS changes that occur in the progression of DN. We describe in detail one such splicing event – the pro- and anti-apoptotic splice isoforms of *BCL-X* (an apoptosis gene). We describe its regulation in a diabetic environment, the association of isoform ratio with DN progression, and its potential as a therapeutic target in DN.

## 2. Materials and Methods

### 2.1 Cell culture and treatments

Conditionally immortalized glomerular endothelial cells (GEnCs) (Satchell et al., 2006) were differentiated in endothelial media (EGM + Bulletkit; Lonza) for 5 days. Treatments were then performed with either glucose soup (GS: 25 mM glucose, 1 ng/ml TNF-α, 1 ng/ml IL-6, and 100 nM insulin), normal glucose (NG: 5.5 mM glucose), or an osmotic control, mannitol (MAN: 5.5 mM glucose and 19.5 mM mannitol), in growth factor deficient EGM media with reduced fetal bovine serum (FBS: 0.5%) (“TREAT” media). Cells were treated daily for 7 days. GEnCs were also treated with varying concentrations of IL-6 (0–50 ng/ml) for 6–24 h in TREAT media. HEK293 cells (ATCC) were sub-cultured from existing cultures in the lab in low glucose DMEM (Sigma-Aldrich) supplemented with 10% FBS or reduced to 0.5% for treatment and 1% pen/strep antibiotics (Sigma-Aldrich). HEK293 cells were treated in either normal glucose (NG) containing 5.5 mM glucose, high glucose (HG) containing 25 mM glucose, diabetic environment treatment named glucose soup plus (GS+) consisting of 25 mM glucose, 1 ng/ml TNF-α, 1 ng/ml IL-6, and 100 nM insulin, mannitol osmotic control (M) also containing 5.5 mM glucose and 19.5 mM mannitol or an alternative diabetic environment medium used to assess the impact of IL-6 named glucose soup minus (GS-) consisting of 25 mM glucose and 1 ng/ml TNF-α. The Oscillating treatment involved changing media from NG to either M, HG, GS+, or GS-back and forth every 12 h till 48 h is reached. Cell lines were genotyped for authentication; Mycoplasma detection is done routinely in our labs.

### 2.2 RNAseq with gene expression and alternative splicing analysis

RNA extraction from GEnCs treated for 1 week with either GS or MAN was performed using the RNAeasy kit (Qiagen). RNA concentration and quality were determined using the TapeStation (Agilent).

High depth and coverage Next Generation Sequencing (NGS) analysis was performed using the Illumina HiSeq 2500 platform (Exeter Sequencing Service). We used 125 bp paired end reads at a high depth and coverage of 125 million reads per sample. Each condition, GS and MAN, had five replicates. Reads were quality-filtered and adapter trimmed using FastQC from the Galaxy bioinformatics platform package. Hisat2, which is able to detect splice junctions, was used to align reads against the reference genome, Hg38. Htseq was used to calculate gene counts before differential analysis using DESeq2. Genes with an FDR (q value) < 0.05 were deemed significantly altered.

MISO (Mixture-of-Isoforms) is a probabilistic framework that was used to quantify the expression level of alternatively spliced genes from our RNAseq data, which was performed by AccuraScience (https://www.accurascience.com/). We implemented a ΔPSI (Percentage Splicing Index) cut-off of 0.2 and Bayes factor cut-off of 10 to select significantly (AS) events between the GS and MAN treated GEnCs.

### 2.3 RT-PCR

RNA was extracted from treated GEnCs using the RNAeasy kit (Qiagen), per the manufacturer instructions. cDNA was synthesized from the RNA using the GoScript™ Reverse Transcription System (Promega). RT-PCR for *BCL-X* splice isoforms was performed using a forward primer positioned in exon 2, before the 5’ splice site for the *BCL-XS* isoform, and a reverse primer positioned in exon 3: *BCL-X* F, 5’-CATGGCAGCAGTAAAGCAAG-3’ and R, 5’-GCATTGTTCCCATAGAGTTCC-3’. The PCR program consisted of 95°C for 120 s, 35 cycles of 95°C for 30 s, 55°C for 30 s and 72°C for 60 s, followed by 72°C for 10 min. A band sized ∼175 bp denoted *BCL-XS* whereas a band sized ∼355 bp denoted *BCL-XL*. A reverse transcriptase negative (RT-) and water control was used for all PCR reactions. PCR products were run on the Biolanalyzer (Agilent) to enable quantification of *BCL-XS* and *BCL-XL* products. The primer sequences for other targets (*ZNF426, ARAP1,* and DOK1) are shown in Supplementary Table 1.

Quantitative RT-PCR (qRT-PCR) was used to assess changes in gene expression using 20 ng cDNA per reaction. The primers are detailed in Supplementary Table 1. SYBR green dye was used with the following PCR conditions using the LightCycler^®^ (Roche): 95°C for 30 s, followed by 40 cycles of 95°C for 1 s and 59°C for 20 s. Relative quantification analysis was performed by normalizing the gene of interest to a housekeeping control (*HPRT1*) following the ΔΔCq method.

### 2.4 Western blotting

Denatured protein samples were run on mini-PROTEAN TGX Stain Free pre-cast gels (4–15%, BIORAD), which allow for visualisation and accurate analysis of the total protein loaded for each sample using a Gel-Doc EZ (BIO-RAD) imaging system. The use of this system means a housekeeping protein loading control is not required as the amount of protein on the membrane for each sample can be quantified. Once the proteins had been transferred on to a polyvinylidene difluoride (PVDF) membrane, total protein could be quantified. Membranes were blocked in 3% bovine serum albumin (BSA) in tris-buffered saline (TBS) plus 0.3% Tween before being probed with anti-BCL-XS/L (Santa Cruz) at 1:1000 dilution in 3% BSA-TBS-Tween (0.3%), at 4°C overnight. After washing membranes in TBS-Tween (0.3%), they were incubated in goat anti-mouse fluorescent secondary antibody (LI-COR) diluted in 3% BSA-TBS-Tween (0.3%), 1:10,000. Membranes were washed again and imaged with the LI-COR Odyssey® CLx. Analysis was performed using the Image Studio software (LI-COR).

### 2.5 Immunofluorescence

GEnCs plated on to coverslips were fixed with 100% methanol for 5 min before blocking with 3% BSA and 5% normal goat serum in phosphate-buffered saline (PBS) for 1 h. Cells were then incubated with anti-SF3B1 (Abcam) at 1:100 dilution in 3% BSA in PBS for 1 h at room temperature. After washing in PBS, the appropriate fluorescent secondary antibody was used (Alexa Fluor) in 3% BSA in PBS for 2 h at room temperature. Sections were then washed in PBS before mounting with gel mount containing DAPI (VECTASHIELD). Images were taken using a Leica DM4000 B LED fluorescent microscope using a 40x objective.

### 2.5 Apoptosis assays

*JC-10 mitochondrial membrane potential assay (Abcam):* This assay detects changes in the mitochondrial membrane potential using the cationic, lipophilic JC-10 dye. In normal cells, JC-10 concentrates in the mitochondrial matrix where it forms red fluorescent aggregates. However, in apoptotic and necrotic cells, JC-10 diffuses out of the mitochondria, changes to monomeric form, and stains cells with green fluorescence. Assays were performed on GEnCs and HEK293 cells plated and treated in a 96-well format in triplicate, per the manufacturers’ protocol.

*Generic caspase activity assay (abcam):* This assay measures the activity of caspase, a widely accepted reliable indicator for cell apoptosis, using TF2-VAD-FMK as a fluorescent indicator that irreversibly binds to activated caspase-1, -3, -4, -5, -6, -7, -8, and -9 in apoptotic cells. Assays were performed on GEnCs plated and treated in a 96-well format in triplicate, per the manufacturers protocol.

*Trypan blue cell viability assay (ThermoFisher Scientific):* Trypan blue stain colors dead cells blue, allowing cell viability to be measured. Assays were performed in GEnCs plated and treated in a 12-well format, per the manufacturers protocol.

### 2.6 Transfections

The transfection was performed on HEK293 cells seeded in a black, clear bottom 96 well culture plate, at a density of 2 x 10^4^ cells per well in a complete growth medium, overnight. The transfection complex comprised 0.2 μg/μl of the plasmid DNA diluted in serum-free medium (Opti-MEM) and gently mixed with Turbofectin 8.0 (Fisher Scientific) at a factor of 3:1. The mixture was incubated for 15 min at room temperature, gently added to the cells, and evenly distributed by carefully rocking.

### 2.7 Human samples and ethics approval

The samples were obtained from participants with and without type 2 diabetes taking part in the BEAt-DKD Exeter and VIBE study in the NIHR Exeter Clinical Research Facility. The samples were only used for purposes identified by the research protocols for which appropriate ethical approval and written consent were obtained. The samples were obtained, processed, and stored following Human Tissue Act 2004. The confidentiality and anonymity of the participants were strictly maintained throughout the study with samples only identified by study ID number and all associated data anonymised. Clinical measurements of diabetes duration, blood pressure, HbA1c, estimated glomerular filtration rate (eGFR), height, weight, urinary albumin/creatinine ratio were obtained.

### 2.8 RNA extraction from blood and urine

*Patient blood samples:* Blood samples (approximately 10 ml) were collected in EDTA-coated tubes and centrifuged for 10 min at 200 x g no more than 4 h after collection. The leukocyte interphase (buffy coat) was collected and placed into TRIzol (ThermoFisher Scientific) before being stored at -80°C for future analysis.

*Patient urine samples:* Patients performed an overnight urine collection at home. The next day, we obtained the urine sample (volumes ranged from 200–3000 ml), which was centrifuged at 2500 x g for 20 min. Urinary sediment (containing the RNA) was resuspended in TRIzol before being stored at -80°C for future analysis.

*RNA extraction:* RNA extraction was performed using the phenol/chloroform method. cDNA synthesis and RT-PCR for *BCL-X* splice isoforms were performed as described above.

### 2.9 Statistical analysis

The data analysis were performed in GraphPad Prism (Prism 10). Unpaired student t-test (parametric) or Mann Whitney U test (nonparametric tests) was used to compare a two group data set and for three or more groups of datasets, a one-way analysis of variance (ANOVA) parametric or Kruskal-Wallis (nonparametric tests) was used. To assess the linear relationship between two sets of data a Pearson or Spearman correlation test (depending on the distribution of the data) was performed following the normality test. All analyses performed were only considered statistically significant if the p value was less than 0.05.

## 3. Results

### 3.1. RNAseq analysis of AS events identifies *BCL-X* splice variants to be associated with DN

In a quest to discover splice variants associated with the progression of DN, we performed an RNAseq analysis on GEnCs. To mimic the diabetic environment, GEnCs were exposed to the GS for 1 week. NG and MAN (to account for osmotic properties) were used as controls. RNA was extracted, short-reads RNAseq analysis was performed, and data were analysed with DESeq2 for gene expression and MISO for differential splice variants.

In the DESeq2 differential expression analysis, using a cut-off of log2 fold change of 0.5 and q<0.05, 124 genes were found to be significantly upregulated and 33 genes significantly downregulated in GEnCs exposed to the glucose soup (GS) for 1 week, in comparison to the mannitol control (**Figure 1A**). Gene ontology (GO) enrichment using the GOrilla tool identified eight GO terms that were significantly enriched (**Figure 1B**). In the heat maps of the two GO terms with the highest enrichment scores, “chemokine receptor binding” and “inflammatory response”, several genes are shown to be involved in diabetes pathogenesis (for example, *CXCL10, IL-6,* and *NFKB1*) (**Figure 1C**). A set of 11 genes (six upregulated and five downregulated) were validated by wet lab analysis (qRT-PCR); nine out of 11 (82%) showed changes in expression as predicted by the bioinformatics analysis (**Figure 1D**).

**Figure 1.**
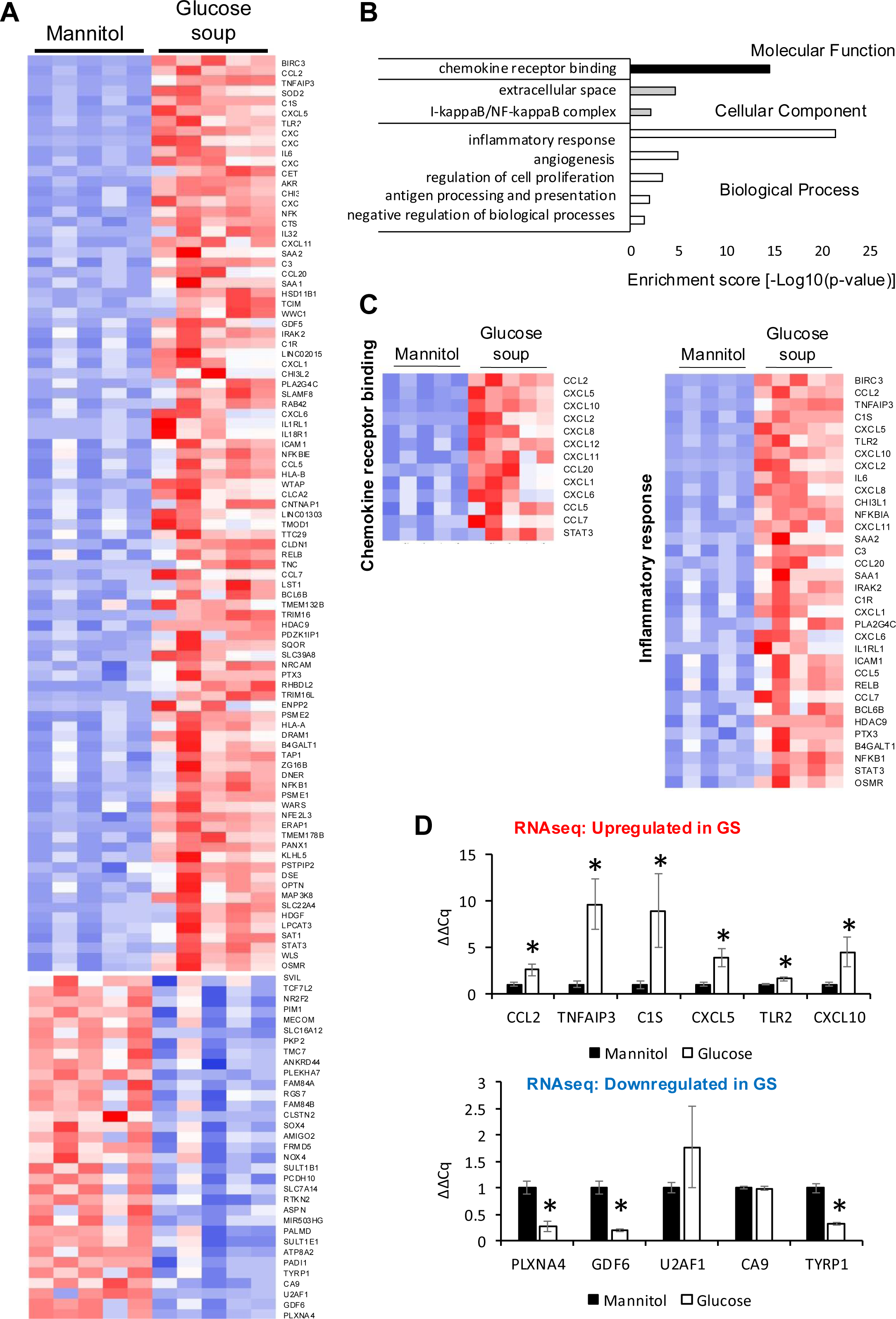
DESeq2 analysis of gene expression changes in glomerular endothelial cells (GEnCs) exposed to a diabetic environment. Differential expression analysis using the DESeq2 Bioconductor package. **(A)** 124 genes were found to be significantly upregulated, and 33 genes significantly downregulated in GEnCs exposed to the glucose soup (GS) for 1 week, in comparison to the mannitol control (n = 5). **(B)** Gene ontology (GO) enrichment using the GOrilla tool identified eight GO terms enriched. **(C)** Heat maps of the two GO terms with the highest enrichment scores, “chemokine receptor binding” and “inflammatory response”. **(D)** Out of 11 genes found to be dysregulated (6 up-regulated and 5 downregulated), 9 were validated using qRT-PCR (82%) (n = 4 biological repeats; *p < 0.05 vs. mannitol control as assessed by Student’s t-test).

MISO (Mixture-of-Isoforms) is a probabilistic framework used to quantify the expression level of alternatively spliced genes from the RNAseq data. We identified 103 significant alternative splicing (AS) events: 5 skipped exon (SE), 12 alternative 5’ splice site (A5SS), 11 alternative 3’ splice site (A3SS), 26 mutually exclusive exon (MXE), and 39 retained intron (RI) events (**Figure 2A**). We implemented a ΔPSI (Percentage Splicing Index) cut-off of 0.2 and Bayes factor cut-off of 10 to select AS events that have changed significantly between conditions. All protein-coding events were validated using RT-PCR and 70% were found to be as predicted by the bioinformatics analysis. Several of the AS events that change upon GS incubation are presented in **Table 1**. Example RT-PCRs of four types of events and the way they change upon GS incubation (*ZNF426, DOK1, ARAP1,* and *BCL-X*) are shown in **Figure 2B**.

**Figure 2.**
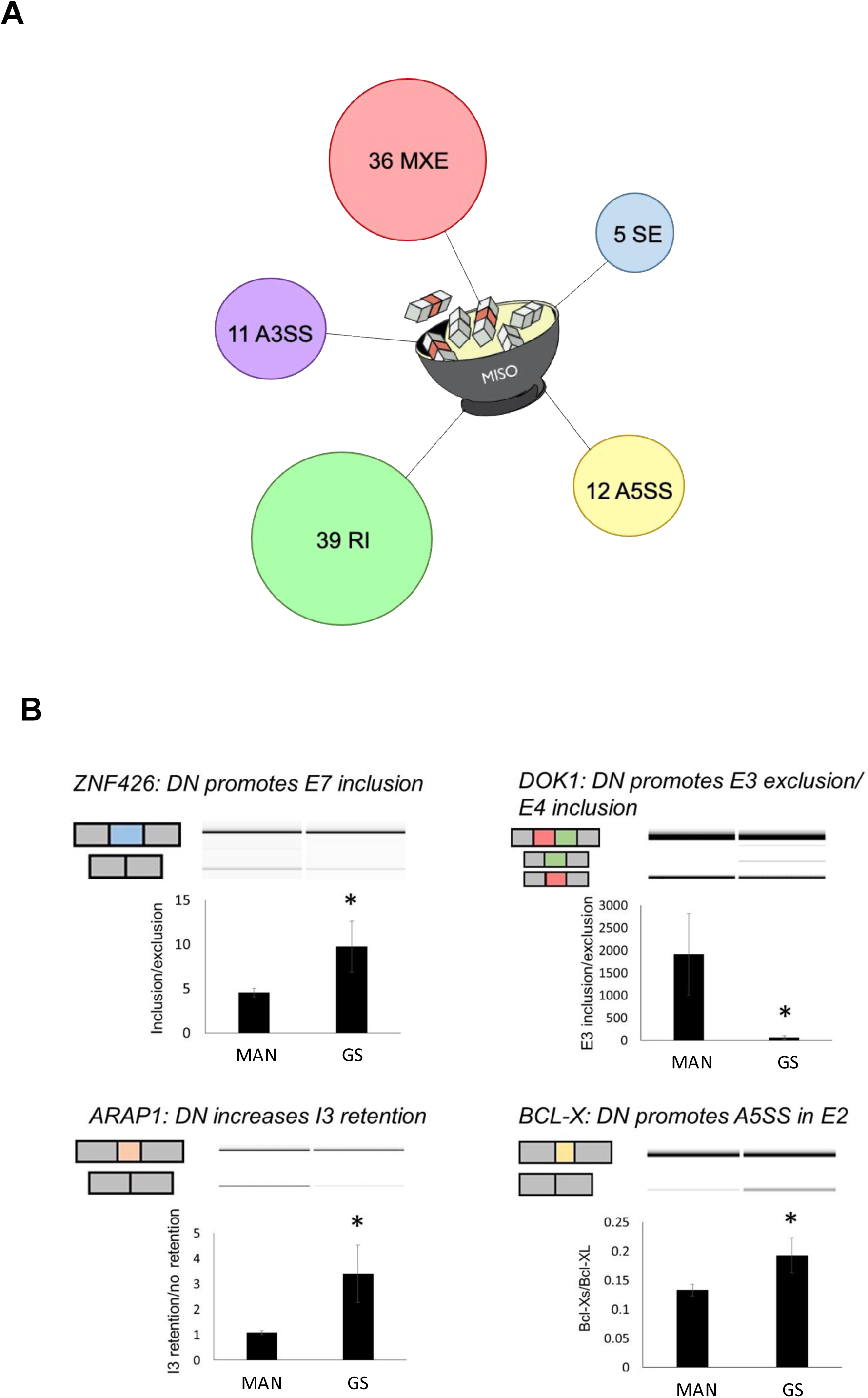
MISO analysis of significant alternative splicing events in glomerular endothelial cells (GEnCs) exposed to glucose soup (GS). MISO (Mixture-of-Isoforms) is a probabilistic framework used to quantify the expression level of alternatively spliced genes from our RNAseq data. (A) Using the MISO framework, we identified 103 significant alternative splicing (AS) events (5 skipped exon (SE), 12 alternative 5’ splice site (A5SS), 11 alternative 3’ splice site (A3SS), 36 mutually exclusive exon (MXE), and 39 retained intron (RI) events). We implemented a ΔPSI (Percentage Splicing Index) cut-off of 0.2 and Bayes factor cut-off of 10 to select significantly (AS) events (n = 5). (B) All events were validated using RT-PCR; an example of four types of events (*ZNF426, DOK1, ARAP1,* and *BCL-X*) are shown (n = 4–6 biological repeats; *p < 0.05 vs. mannitol (MAN) control as assessed by Student’s t-test). We were able to validate 70% of protein-coding AS events.

The list of splicing events that move when exposed to a diabetic environment, as described in **Table 1**, included a switch in *BCL-X* isoforms. As described above, *BCL-X* has two splice isoforms, *BCL-XS* and *BCL-XL,* which result from a 5’ alternative splice event (**Figure 3A**) and have opposing functions on apoptosis (10). The functional importance of this splicing switch has been studied extensively in cancer (11); therefore. we wanted to explore its involvement in the pathogenesis of DN.

**Figure 3.**
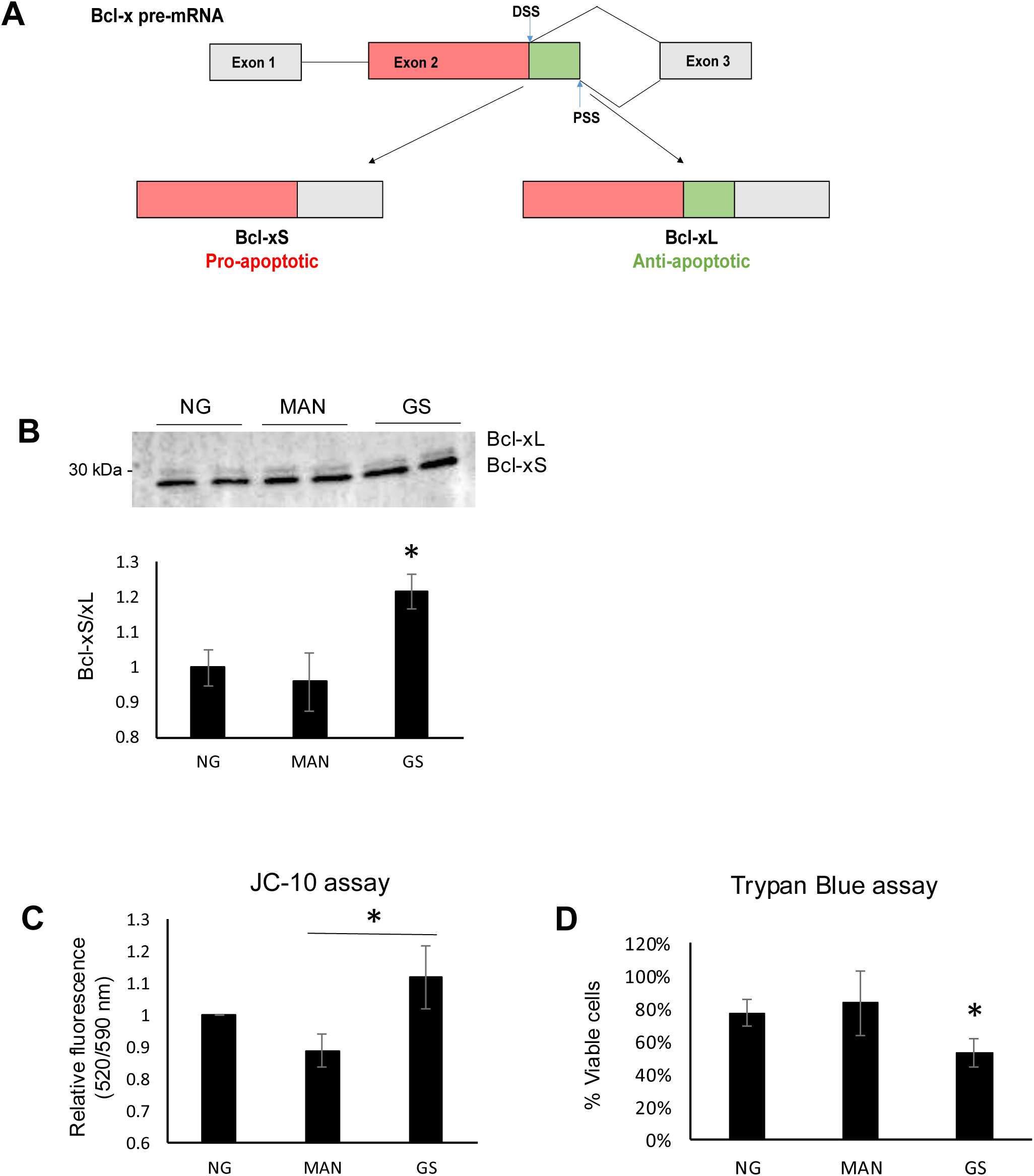
Glomerular endothelial cells (GEnCs) exposed to a diabetic environment have an upregulation of *BCL-XS/BCL-XL*, resulting in apoptosis. (A) Diagrammatic representation of the *BCL-X* alternative splicing (AS) event. Use of an alternative 5’ splice site in exon 2 of the *BCL-X* pre-mRNA results in the expression of the pro-apoptotic *BCL-XS* isoform, in contrast to the anti-apoptotic *BCL-XL*. (B) The switch in *BCL-X* AS to promote BCL-XS expression in response to treatment of GEnCs with glucose soup (GS) was confirmed at the protein level using Western blotting (n = 4 biological repeats; *p < 0.05 vs. mannitol (MAN) and normal glucose (NG) controls as assessed by one-way ANOVA and Tukey post-hoc test). (C) GEnCs treated with GS showed an increase in the fluorescence green/red (520/590 nm) ratio, representing an increase in cell apoptosis, compared to MAN control (n = 4 biological repeats; *p < 0.05 vs. MAN as assessed by one-way ANOVA and Tukey post-hoc test). (D) GEnCs treated with GS also had a significantly reduction in viability (n = 3 biological repeats; *p < 0.05 vs. mannitol (MAN) and normal glucose (NG) controls as assessed by one-way ANOVA and Tukey post-hoc test).

### 3.2. The diabetic environment increases the *BCL-XS/BCL-XL* ratio in GEnCs and HEK293 cells, increasing apoptosis

The analysis of *BCL-X* AS described above was performed at the RNA level; however, the ultimate effectors of cellular function are proteins. Therefore, we investigated whether BCL-X protein isoforms also change as expected. As observed in the western blotting analysis in **Figure 3B**, upon exposure of GEnCs to GS, the ratio of BCL-XS to BCL-XL is increased. The same ratio does not increase upon exposure to mannitol. Furthermore, we confirmed that this switch in isoforms functionally results in an increase in apoptosis and cell death using the JC-10 and Trypan Blue assay (**Figures 3C** and **3D**).

We wanted to further explore the diabetic environment in which high-glucose induces the BCL-X splicing switch and apoptosis. Therefore, we tested other conditions, including incubation of cells with only high-glucose (HG) instead of GS, or changes in the exposure mode such as oscillating high-glucose (OHG)/oscillating glucose soup (OGS). Oscillating treatment involves changing media from normal glucose to either MAN or HG back and forth every 12 h, for 48 h. As observed in **Supplementary Figures 1–4** OHG, GS, and OGS, but not HG, increased the ratio of BCL-XS to BCL-XL, as well as apoptosis, in HEK293 cells.

### 3.3. Overexpression of the *BCL-XL* isoform rescues the apoptosis induced by the diabetic environment

There are many determinants of apoptosis that are active during the progression of DN. In addition, the *BCL-X* splicing switch might be a consequence of the DN development. Therefore, the functional importance of this splicing switch needs to be investigated, i.e., if we switch back the isoform ratio to that of the physiologically normal situation by increasing the expression of anti-apoptotic *BCL-XL*, can we see a rescue, at least partially, of apoptosis?

To address this question, we used plasmid-induced overexpression of the anti-apoptotic *BCL-XL* isoform (see **Supplementary Figure 5** for proof of expression). HEK293 cells, with or without *BCL-XL* overexpression, were exposed to NG, MAN, and GS. As seen in **Figure 4**, there is a significant decrease in apoptosis induced by GS exposure in cells transfected with the *BCL-XL* plasmid.

**Figure 4.**
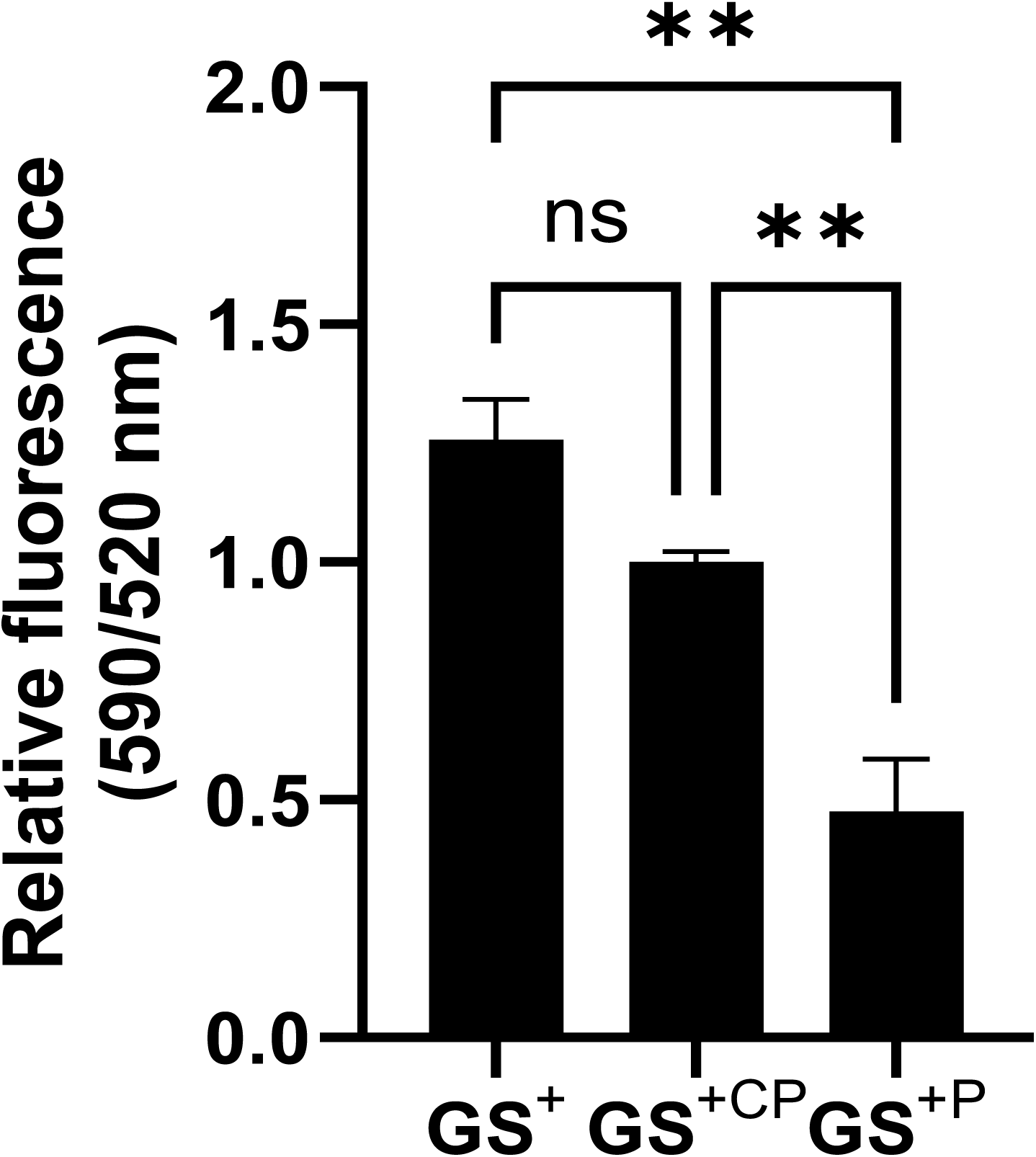
Transient overexpression of *BCL-XL* prevents HEK293 cell apoptosis. HEK293 cells were plated and incubated for 24 h then treated with Glucose Soup (25 mM with TNFα 1 ng/ml, IL-6 1 ng/ml and insulin 100 nM) GS^+^. Cells were transfected without plasmid or with a control plasmid (CP) or BCL-XL plasmid (P) at 0.2 μg using turbofectin 24 h after treatment. The JC-10 apoptosis assay was performed 48 h after treatment (the 520/590nm fluorescence ratio is proportional to the amount of apoptosis). Statistical analysis, performed using an ordinary one-way ANOVA and a multiple comparison analysis, showed a significant decrease between GS^CP^ and GS^+P^ (mean = 0.48 ± 0.11 SEM, **p=0.007, n = 3 repeats), but not between GS^CP^ and GS^+^ (mean = 1.26 ± 0.09 SEM, p=0.11, n = 3 repeats).

### 3.4. The *BCL-XS/BCL-XL* ratio correlates with various clinical parameters measured in the blood and/or urine of patients with DN

The *in vitro* studies showed that the *BCL-X* splicing isoform ratio changes in renal cells exposed to a diabetic environment resulted in an increase in the *BCL-XS* isoform and, subsequently, an increase in cellular apoptosis. Moreover, overexpression of the *BCL-XL* isoform can partially rescue the apoptosis. This suggests that the *BCL-X* splicing switch is important in the development of the DN. Therefore, we investigated whether the *BCL-X* isoform ratio is associated with clinical parameters of the progression of DN.

We analyzed 45 blood samples and 28 urine samples from healthy and diabetic patients with varying stages of nephropathy. Patients were enrolled in the Biomarker Enterprise to Attack Diabetic Kidney Disease (BEAt-DKD) study(12), an EU IMI funded study which is a partnership between public and private sectors which aims to improve prevention and management of diabetic kidney disease, and the VIBE study (see acknowledgements), including patients with type 2 diabetes (see Table 2). RNA was extracted from the urinary sediment or leukocytes, RT-PCR performed for the *BCL-X* splice isoforms (see an example **Supplementary Figure 6**), and the ratio of isoforms was found to correlate with various clinical parameters.

A negative correlation was observed between the estimated glomerular filtration rate (eGFR) and the *BCL-XS/BCL-XL* ratio from urine sediments (Pearson r = -0.3934, p = 0.0468); however, eGFR did not correlate with the *BCL-XS/BCL-XL* ratio in leucocytes (Spearman r = -0.04143, p = 0.972) (**Figure 5A** and **B**). The urinary albumin creatinine ratio (UACR) correlated positively with the leucocyte *BCL-XS/BCL-XL* ratio (Pearson r = 0.4522, p = 0.0038), but it did not correlate with *BCL-XS/BCL-XL* ratio in urinary sediment (Spearman r = -0.05321, p = 0.8141) (**Figure 5C** and **D**). Microalbuminuric patients did not show a change in the *BCL-XS/BCL-XL* expression ratio in leukocytes; however, the *BCL-XS/BCL-XL* expression ratio obtained from urine showed a significant increase in microalbuminuric patients compared to controls but not compared to normoalbumuric participants with diabetes (**Figure 5E** and **F**). Other parameters, including serum creatinine, Hb1Ac, fasting glucose, and gender, did not correlate with the ratio of *BCL-X* isoforms ratio (**Supplementary Figures 7-10**). The fact that the *BCL-X* isoform ratio correlated with some of the parameters of DN progression further strengthens the conclusion from the *in vitro* studies that this splicing event is important in development of DN.

**Figure 5.**
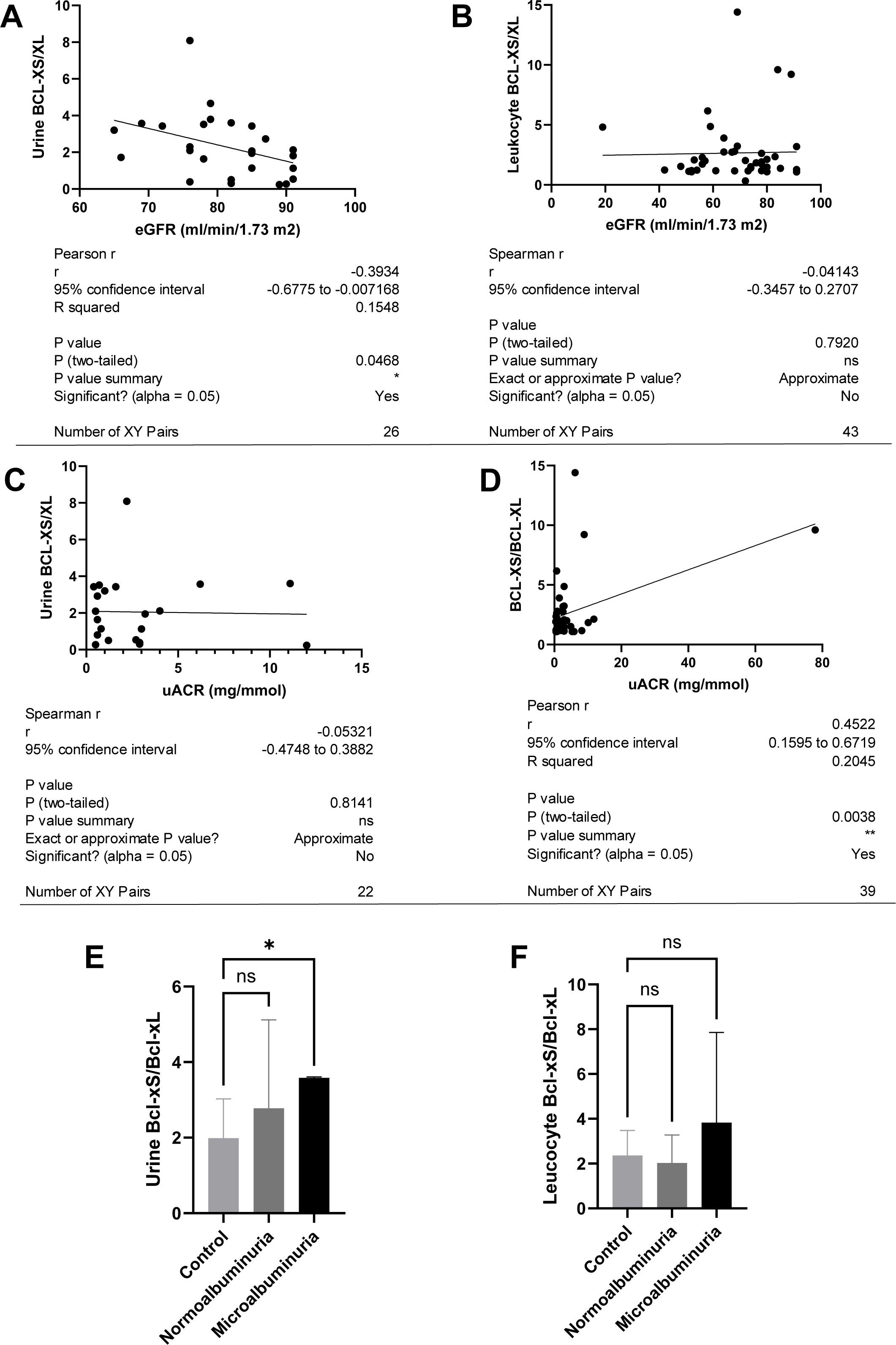
The *BCL-X* isoform ratio correlates with some clinical parameters in diabetic nephropathy patients. **(A)** The *BCL-XS/BCL-XL* ratio obtained from urine sediment has a negative correlation with the estimated glomerular filtration rate (eGFR): cDNA was generated from RNA extracted from sediments of urine collected from patients overnight. RT-PCR was performed using *BCL-X* primers to amplify the long and short isoforms. The bioanalyser was used to quantify the isoform concentration and the *BCL-XS/BCL-XL* ratio was calculated. A plot of eGFR against the urinary sediment *BCL-XS/BCL-XL* ratio was generated using GraphPad prism. The data was analysed using Pearson correlation analysis (r = 0.3934, p = 0.0468, n = 26 overnight urine samples). **(B)** The *BCL-XS/BCL-XL* ratio obtained from leucocytes does not correlate with the eGFR: cDNA was generated from RNA extracted from leucocytes. A Spearman correlation analysis was performed for the data (r = -0.04143, p = 0.7920, n = 43 blood samples). **(C)** The *BCL-XS/BCL-XL* ratio from urine sediment does not correlate with the urine albumin creatinine ratio (uACR) (Spearman r = -0.05321, ns p = 0.9187, n = 22 samples). **(D)** The *BCL-XS/BCL-XL* ratio from leucocytes has a positive correlation with the uACR in participants without clinically significant albuminuria (Pearson r = 0.4522, p = 0.0361, n = 39 blood samples). **(E)** The *BCL-XS/BCL-XL* ratio obtained from urine sediment is not significantly increased in normoalbuminuria but significantly increased in microalbuminuria compared to control. The data was statistically analysed us a Mann– Whitney test. Normoalbuminuria (mean = 2.78; ns p = 0.507; n = 11 patients; SEM ± 0.7049); Microalbuminuria (mean = 3.584; *p = 0.0147; n = 2 patients; SEM ± 0.0159). **(F)** The leucocyte *BCL-XS/BCL-XL* ratio is not significantly increased in both normoalbuminuria and microalbuminuria compared to control. The data was statistically analysed by a Man–Whitney test. Normoalbumiuria (mean = 2.03; ns p = 0.352; n = 24 patients; SEM ± 0.255); Microalbuminuria (mean = 3.83; ns p > 0.999; n = 15 patients; SEM ± 1.04).

### 3.5. An increase in *IL-6* regulates *BCL-X* splicing to promote *BCL-XS* expression in diabetic GEnCs

The experiment described above highlighted the importance of *BCL-X* splicing in DN; therefore, we explored which molecules are important for its regulation in this context.

IL-6 has previously been reported to repress *BCL-XL* expression through an intron retention element (IRE) in intron 2 of the *BCL-X* pre-mRNA (10), although the exact mechanism is unknown (**Figure 6A**). Analysis of RNAseq data described above showed that *IL-6* expression is increased in GEnCs exposed to a diabetic environment (GS treatment) (**Figure 6B**). Indeed, qRT-PCR validation showed a clear increase in *IL-6* expression upon exposure to GenC. The GS contains *IL-6*; however, treating cells with GS that did not contain IL-6 showed a similar increase in IL-6 expression (**Figure 6C**). Treatment of GEnCs with IL-6 resulted in a dose-dependent increase in the ratio of *BCL-XS* to *BCL-XL* (**Figure 6D**), in addition to a corresponding increase in apoptosis as ascertained by JC-10 and caspase activation assays (**Figure 6E** and **F**), and a decrease in viability shown with the Trypan Blue assay (**Figure 6G**). Furthermore, exposure of HEK293 cells to increasing concentrations of IL-6 resulted in a dose-dependent increase in the ratio of *BCL-XS* to *BCL-XL* and a corresponding increase apoptosis (**Supplementary Figure 11)**.

**Figure 6.**
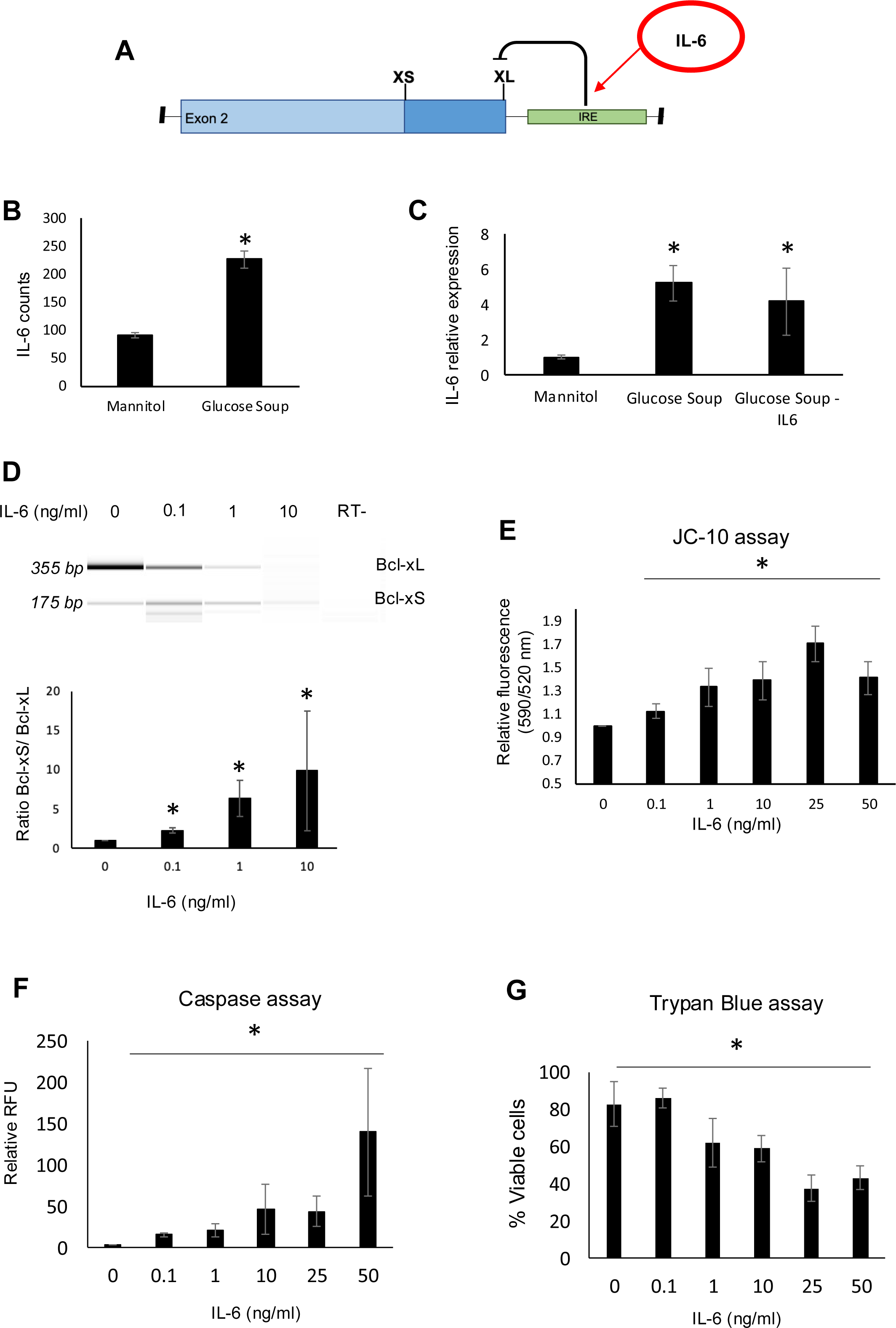
IL-6 is a major regulator of *BCL-X* splicing. **(A)** Schematic of *BCL-X* exon 2 and position of intron response element (IRE). **(B)** RNAseq and DESeq2 analysis of GEnCs treated with glucose soup (GS) showed a significant increase in the IL-6 read counts compared to mannitol (MAN) control (n = 5; *p < 0.05). **(C)** The increase in IL-6 expression in response to GS was confirmed with qRT-PCR, which persisted in the absence of IL-6 from the GS treatment of GEnCs (n = 3; *p < 0.05 vs. mannitol control as assessed by one-way ANOVA and Tukey post-hoc test). **(D)** Treatment of GEnCs for 6 h with increasing concentrations of IL-6 alone resulted in a dose-dependent increase in *BCL-XS/BCL-XL* (n = 3; *p < 0.05 vs. 0 ng/ml IL-6 as assessed by one-way ANOVA and Tukey post-hoc test). **(E)** IL-6 dose-dependently increased the fluorescence green/red (520/590 nm) ratio, representing an increase in cell apoptosis, using the JC-10 assay (n = 4; *p < 0.05 vs. 0 ng/ml IL-6 as assessed by one-way ANOVA and Tukey post-hoc test). **(F)** IL-6 dose-dependently increased caspase activation (n = 4; *p < 0.05 vs. 0 ng/ml IL-6 as assessed by one-way ANOVA and Tukey post-hoc test). **(G)** IL-6 dose-dependently decreased cell viability, as assessed with Trypan blue staining, indicating increased cell apoptosis with increasing IL-6 (n = 4; *p < 0.05 as assessed by one-way ANOVA).

### 3.6. Expression of splice factors *SF3B1* and *PTBP1* is decreased in a diabetic environment, which may regulate the *BCL-X* splicing switch

We further investigated which splice factors may be involved in the regulation of *BCL-X* splicing in the diabetic context.

SF3B1 has previously been reported to promote *BCL-XS* splice site selection through binding to ceramide-responsive RNA cis-element 1 (CRCE1) in exon 2 of the *BCL-X* pre-mRNA (10) (**Figure 7A**). RNAseq and DESeq2 analysis of GEnCs treated with GS showed a significant decrease in the *SF3B1* read counts compared to MAN control; this was validated by qRT-PCR (**Figure 7B**). The downregulation of SF3B1 when GEnCs are incubated in GS was confirmed at the protein level with immunofluorescence (**Figure 7C**). Interestingly, the expression of SF3B1 is not regulated by IL-6 (**Supplementary Figure 12**).

**Figure 7.**
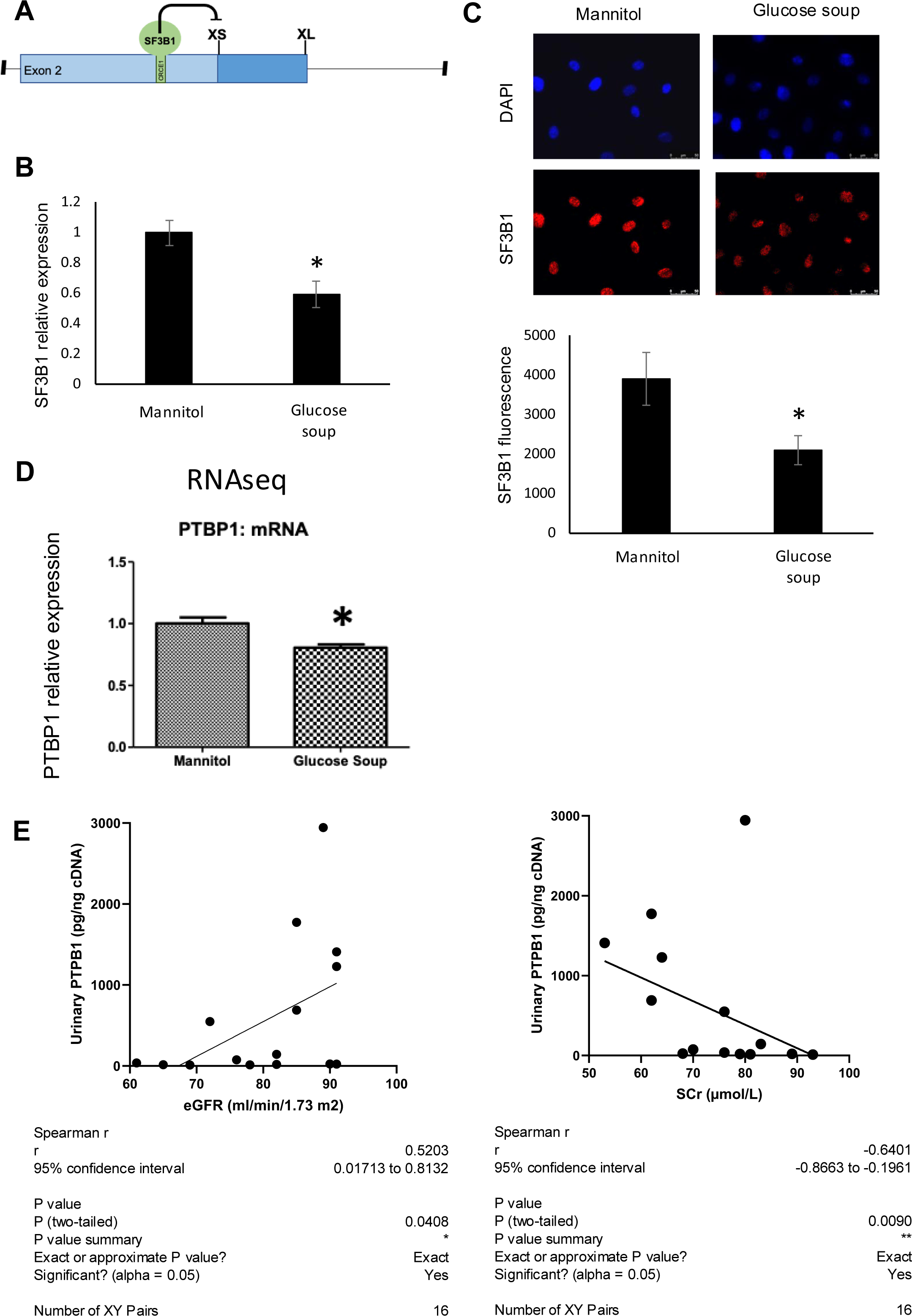
The splice factors *SF3B1* and *PTBP1* are downregulated in response to glucose soup (GS) treatment of glomerular endothelial cells (GEnCs). **(A)** *SF3B1* binds to a ceramide-responsive RNA cis-element 1 (CRCE1) in exon 2 of the *BCL-X* pre-mRNA and represses *BCL-XS*. **(B)** qRT-PCR shows decrease in *SF3B1* expression when cells are incubated in GS (n = 4; *p < 0.05 vs. mannitol (MAN) control, as assessed by Student’s t-test). **(C)** Quantitation of immunofluorescence for SF3B1 confirms the decrease at protein level (n = 3; *p < 0.05 vs. MAN control, as assessed by Student’s t-test). **(D)** RNAseq data showing a decrease in *PTBP1* in GEnCs treated with GS. **(E)** Urinary *PTBP1* expression correlated with the estimated glomerular filtration rate (eGFR) and serum creatinine (SCr) in the urine samples from diabetic patients (eGFR: Spearman r = 0.5203, p = 0.0408, n = 16; SCr: Spearman r = -0.6401, p = 0.009, n = 16).

Another splice factor reported to regulate *BCL-X* splicing, *PTBP1*, is decreased when GEnCs are incubated in GS (based on RNAseq data) (**Figure 7D**). Interestingly, the urinary *PTBP1* expression in patients with various degrees of DN correlated with markers of a decline in renal function, i.e., a positive correlation with the eGFR (Spearman r = 0.5203; p = 0.0408) and a negative correlation with serum creatinine (Spearman r = -0.6401; p = 0.009) (**Figure 7E**).

## Discussion and Conclusions

Alternative splicing (AS) is a major level of gene regulation that is able to drive various functions in the cells sometimes autonomously from transcription. Splicing variants are described in virtually every class of molecules from growth factors to tyrosine kinase/phosphatase receptors to tumour suppressors and oncogenes. Many times, the splicing isoforms have opposing functions e.g., pro- or anti-angiogenic, pro- or anti-apoptotic (7, 13). There is recent evidence that the occurrence of a particular set of splice isoforms is determinant for both physiological and pathological processes (14). Indeed, there are quite a few examples were various biological processes are coordinated by networks of alternative splicing regulators - e.g. tissue and organ development (15, 16), cholesterol biosynthesis and uptake (17), pancreatic beta cells function and survival (18) or epithelial-mesenchymal transitions (19). Interestingly, a large number of apoptosis gene’s function is controlled by AS (20, 21). It is highly likely that this may occur in other processes, including DN pathogenesis and may represent a novel paradigm in biology.

Diabetes is characterized by hyperglycemia and the activation of proinflammatory cytokines. Hyperglycemia has been shown to cause programmed cell death in a variety of renal cells including endothelial and podocyte cells which has been implicated in the pathogenesis of DN (22, 23). Therefore, understanding the mechanism involved in the anti-apoptotic to pro-apoptotic splicing in Bcl-x (a major apoptosis gene) in renal cells may be a key to rescuing renal cells from apoptosis and could also serve as a novel biomarker for DN severity.

This study aimed to determine whether alternative splicing events identified in DN can be used in a therapeutic strategy in DN as well as whether they are associated with DN severity.

We discovered in this study that the Bcl-x AS event is dysregulated in diabetic GEnC by RNAseq analysis. Recognizing the associated programmed cell death that comes with diabetic nephropathy, we sought to further understand the mechanism of the Bcl-x gene AS in diabetic conditions and how it can be exploited to rescue diabetic cells from apoptosis as well as utilized in the development of a biomarker.

Systemic increase in IL-6 together with other cytokines has been associated with obesity-related inflammation. This increase has also been implicated as a risk factor in the development of insulin resistance as well as T2D (24, 25). Studies have shown that IL-6 regulates Bcl-x gene AS in K562 cells in favour of the pro-apoptotic Bcl-xS isoform (26). RT-PCR for the Bcl-x isoforms showed an increase in Bcl-xS band intensity as IL-6 concentration increased from 0.1 ng/ml to 50 ng/ml. Moreover, the JC-10 assay showed an increase in apoptosis of HEK293 as IL-6 concentration increased. This indicates that Bcl-x AS may be dependent on IL-6 in a dose-wise manner.

Hyperglycemia is a hallmark of diabetes and has also been associated with high IL-6 levels(27). Consequently, it was necessary to determine the impact of a high glucose environment on Bcl-x AS. GS+ treatment caused a significant increase in the Bcl-xS/Bcl-xL ratio compared to NG as well as a significant increase in apoptosis of HEK293 cells in GS+ treatment compared to NG using a JC-10 assay.

Additionally, studies have demonstrated that oscillating high glucose could be more deleterious on endothelial function in comparison to stable or continuous high glucose as well as hyper activation of p53, a molecule known to be involved in apoptosis(28). Investigating the effects of oscillating high glucose (OHG) on Bcl-x AS showed that OHG treatment caused a significant increase in Bcl-xS/Bcl-xL ratio compared to NG with a corresponding increase in apoptosis of HEK293 cells in OHG treatment compared to NG treatment using the JC-10 assay. Oscillating glucose soup (OGS+) treatment also showed a significant increase in Bcl-xS/Bcl-xL ratio compared to NG. The JC-10 results showed a significant increase in apoptosis of HEK293 cells treated with OGS+ compared to NG treatment. We therefore concluded that interleukin-6 or a combination of oscillating high glucose and interleukin-6 play a role in Bcl-x gene AS, which consequently leads to apoptosis activation

We saw a significant decrease in apoptosis between HEK293 cells treated in GS+ and then transfected with Bcl-xL transiently. This was compared to HEK293 cells treated with GS+ with transient transfection of a control plasmid. However, there was no significant difference between HEK293 cells treated in GS+ with no transfection compared to HEK293 cells transiently transfected with a control plasmid. This preliminary work serves as evidence that there is a possibility that cells can be protected from apoptosis resulting from hyperglycemia through Bcl-xL upregulation.

Using Pearson correlation analysis, we attempted to establish a relationship between kidney function parameters in research participants and urine sediment or leucocyte Bcl-xS/Bcl-xL ratio. The kidney function metric eGFR was discovered to weakly correlate with the Bcl-xS/Bcl-xL ratio acquired from patients’ urine sediments, whilst uACR moderately correlated with the Bcl-xS/Bcl-xL ratio obtained from leucocytes from patient blood. It is important to note that, while serum creatinine is used to assess renal function, additional confounding factors influence creatinine production and secretion. Muscle mass is a factor that influences creatine synthesis, levels, and, consequently, creatinine production. Age and gender appear to be the most important factors in influencing total muscle mass and, thus, creatinine production. Men create more creatinine than women because they have larger muscular mass. Lower creatinine production can also result from age-related muscle mass reduction (29). Furthermore, the amount of dietary creatine taken, typically in the form of meat, influences creatinine synthesis as well as total muscle mass (30). Reduced creatinine synthesis is associated with pathophysiologic diseases that cause muscle atrophy and decrease muscle mass. These circumstances include long-term glucocorticoid medication, muscular dystrophy, and paralysis just to mention a few. Since multiple unconnected factors may have simultaneous effects on serum creatinine levels, it is not ideal for renal function assessment. However, it is much more advantageous to use serum creatinine, due to its low cost and availability. These drawbacks in serum creatinine levels estimation could be the reason Bcl-xS/Bcl-xL ratio from leucocyte and urine sediment do not correlate these parameters. Despite the limits and inconveniences of measuring albuminuria, chronic disease markers have been employed for many years. However, in the lack of other known and tested approaches, they are used continuously with corrections and considerations, making them more onerous. Other urine markers mentioned in reviews include Neutrophil gelatinase-associated lipocalin (NGAL), N-acetyl-beta-glucosaminidase (NAG), type IV collagen, and nephrin, to name a few (31). These markers are expected to be more durable than those now available. Future research could investigate the link between these new indicators and Bcl-x AS. Furthermore, a larger cohort will solidify our findings.

In conclusion, we discovered that oscillating high glucose and interleukine-6 impact Bcl-x gene AS, increasing the pro-apoptotic isoform Bcl-xS and inducing apoptosis in HEK293 cells. We further propose that increasing the anti-apoptotic isoform Bcl-xL protects HEK293 cells from apoptosis. Finally, the Bcl-xS/Bcl-xL ratio correlates with renal function measures, eGFR and uACR, and may serve as a potential predictive tool for DN severity.

## Data availability

The data that support the findings of this study are available from the corresponding author upon reasonable request.

## Acknowledgements

The BEAt-DKD project has received funding from the Innovative Medicines Initiative 2 Joint Undertaking (JU) under grant agreement No 115974 (BEAt-DKD). This Joint Undertaking receives support from the European Union’s Horizon 2020 research and innovation programme and EFPIA with JDRF. The VIBE study was funded by Diabetes UK (grant reference: 16/0005489). The study was also supported by the National Institute for Health Research (NIHR) Exeter Clinical Research Facility. The views expressed are those of the author(s) and not necessarily those of the NIHR or the Department of Health and Social Care.

Many thanks to Kate Bosson for helping with some of the experiments.

## Contribution statement

Megan Stevens - conception and design, acquisition of data, analysis and interpretation of data, drafting the article, final approval

Monica Ayine - acquisition of data, analysis and interpretation of data, drafting the article, final approval

Kim Gooding - conception and design, reviewing the article, final approval

Angela Shore - conception and design, reviewing the article, final approval

Pedro Marqueti – acquisition of data, reviewing the article, final approval

Sebastian Oltean - conception and design, analysis and interpretation of data, drafting the article, final approval

## Supplementary Figures

**Supplementary Figure 1. Oscillating high glucose significantly increases Bcl-xS/Bcl-xL ratio.**

RT-PCR samples from HEK293 cells treated with Normal glucose (5.5 mM) **NG**, Mannitol **M**, Oscillating Mannitol **OM**, High Glucose (25mM) **HG** or Oscillating high glucose **OHG** for 48 hrs were run on the bioanalyzer. Oscillating treatment involves changing media from **NG** to either **M** or **HG** back and forth every 12 hrs till 48 hrs is reached **A**. bioanalyzer gel image. **B**. quantification of bioanalyzer concentrations showed a significant increase between **NG** and **OHG** (mean=2.04, ±0.28 SEM, ****p<0.0001, N=4 repeats) and between **OM** and **OHG** (mean=1.10, ±0.05 SEM, ***p=0.0005, N=4 repeats). Data were analysed using ordinary one-way ANOVA with a multiple comparison post hoc analysis.

**Supplementary Figure 2. Oscillating high glucose treatment increases apoptosis in HEK293 cells**

HEK293 cells were treated with Normal glucose NG (5.5mM), Mannitol M, Oscillating Mannitol OM, High Glucose (25mM) HG and Oscillating High Glucose OHG for 48 hrs. A JC-10 assay was performed after the 48hours to assess apoptosis. The graph represents a ratio of apoptotic cells (green) to normal cells (red). Statistical analysis was performed using an ordinary one-way ANOVA. Statistically significant increase in apoptosis for the pairwise analysis between NG and OHG (mean=1.30, ±0.061 SEM, *p=0.031, N=10 repeats), HG and OHG (mean=1.30, ±0.061 SEM, **p=0.037, N=10 repeats) and OM and OHG (mean=1.30, ±0.061 SEM, **p=0.047, N=10 repeats).

**Supplementary Figure 3. Glucose soup, but not oscillating glucose soup significantly increases Bcl-xS/Bcl-xL ratio.**

RT-PCR samples from HEK293 cells treated with Normal glucose (5.5 mM) NG, Mannitol M, Oscillating Mannitol OM, Glucose Soup (25 mM with TNFα 1ng/ml, IL-6 1ng/ml and insulin 100 nM ) GS+ or Oscillating glucose soup OGS+ for 48 hrs were run on the bioanalyzer. Oscillating treatment involves changing media from NG to either M or GS+ back and forth every 12 hrs till 48 hrs is reached A. bioanalyzer gel image. B. quantification of bioanalyzer concentrations showed a significant increase between NG and GS+ (mean=1.89, ±0.28 SEM, *p<0.025, N=4 repeats) and between M and GS+ (mean=1.01, ±0.13 SEM, *p=0.027, N=4 repeats). Data were analysed using ordinary one-way ANOVA with a multiple comparison post hoc analysis.

**Supplementary Figure 4. Glucose soup but not oscillating glucose soup treatment increase apoptosis in HEK293 cells.**

HEK293 cells were treated with Normal glucose NG (5.5mM), Mannitol M, Oscillating Mannitol OM, Glucose soup (25mM) GS+ and Oscillating Glucose Soup OGS+ for 48 hrs. A JC-10 assay was performed after the 48hours to assess apoptosis. The grapth represents a ratio of apoptotic cells (green) to normal cells (red). Statistical analysis was performed using an ordinary one-way ANOVA. A statistically significant increase in apoptosis was seen in the pairwise analysis between NG and GS+ (mean=2.98, ±0.33 SEM, ****p<0.0001, N=17 repeats), M and GS+ (mean=2.98, ±0.33 SEM, ****p<0.0001, N=17 repeats) as well as between GS+ and OGS+ (mean=2.98, ±0.33 SEM, **p=0.005, N=17 repeats).

**Supplementary Figure 5. Transient transfection of HEK293 cells**. Cells were transfected with a Bcl-xL plasmid expressing EGFP using turbofectin transfection kit for 48 hours and treated with ampicillin (100 mg/ml). Images taken with EVOS-FL, 10X magnification

**Supplementary Figure 6. Example of RT-PCR analysis of Bcl-x splice isoforms in patients urine samples**

RNA has been isolated from urine sediments and RT-PCR with specific primers for the two Bcl-x splice isoforms has been performed

**Supplementary Figure 7. A. Bcl-xS/Bcl-xL ratio from urine sediment does not correlate with serum creatinine ratio**: cDNA was obtained from RNA extracted from sediments of urine collected from patients overnight. RT-PCR was performed using Bcl-x primers to amplify the long and short isoforms. The bioanalyser was used to quantify isoform concentration and Bcl-xS/Bcl-xL ratio was calculated. A graph of the Serum creatinine against urinary sediment Bcl-xS/Bcl-xL ratio was generated using GraphPad prism. The data were analysed with Pearson correlation analysis followed by linear regression analysis. (r=0.257, ns p=0.188, N=28 overnight urine samples)

**B. Bcl-xS/Bcl-xL ratio obtained from leucocytes has no correlation with serum creatinine ratio**: cDNA was obtained from RNA extracted from leucocytes. RTPCR was performed using Bclx primers to amplify the long and short isoforms. The bioanalyser was used to quantify isoform concentration and BclxS/BclxL ratio was calculated. A graph of Scr against leucocyte Bcl-xS/Bcl-xL ratio was generated using GraphPad prism. A Pearson correlation analysis was performed for the data set, including linear regression analysis. (r=0.010, ns p=0.948, N=43 blood samples). Log transformation was used to trans form the data before Pearson correlation was applied.

**Supplementary Figure 8.**

**A.** Bclxs/BclxL ratio from urine sediments does not correlate with glycated haemoglobin: cDNA was obtained from RNA extracted from sediments of urine collected from patients overnight. RTPCR was performed using Bclx primers to amplify the long and short isoforms. The bioanalyser was used to quantify isoform concentration and Bclxs/Bclxl ratio was calculated. A graph of HbA1c against urinary sediment Bcl-xS/Bcl-xL ratio was generated using GraphPad prism. A Pearson correlation analysis and a linear regression analysis were performed for the data set. (r=-0.065, ns p=0.74, N=28 overnight urine samples)

**B.** Bclxs/BclxL ratio obtained from leucocytes has no correlation with glycated haemoglobin: cDNA was obtained from RNA extracted from leucocytes. RTPCR was performed using Bclx primers to amplify the long and short isoforms. The bioanalyser was used to quantify isoform concentration and Bclxs/Bclxl ratio was calculated. A plot of HbA1c against leucocyte Bcl-xS/Bcl-xL ratio was generated using GraphPad prism. Data were analysed using Pearson correlation analysis and a linear regression analysis was also performed. (r=-0.115, ns p=0.464, N=43 blood samples)

**Supplementary Figure 9.**

**A.** Bclxs/BclxL ratio from urine sediment does not correlate with Fasting Glucose: cDNA was obtained from RNA extracted from sediments of urine collected from patients overnight. RTPCR was performed using Bcl-x primers to amplify the long and short isoforms. The bioanalyser was used to quantify isoform concentration and Bclxs/Bclxl ratio was calculated. A plot of Fasting Glucose against urinary sediment Bcl-xS/Bcl-xL ratio was generated using GraphPad prism. A Pearson correlation was performed as well as a linear regression analysis (r=-0.220, ns p=0.292, N=25 overnight urine samples)

**B.** Bclxs/BclxL ratio obtained from leucocytes does not correlate with Fasting Glucose rate: cDNA was obtained from RNA extracted from leucocytes. RTPCR was performed using Bcl-x primers to amplify the long and short isoforms. The bioanalyser was used to quantify isoform concentration and BclxS/BclxL ratio was calculated. A plot of Fasting Glucose against leucocyte Bcl-xS/Bcl-xL ratio was generated using GraphPad prism. A Pearson correlation analysis was performed for the data as well as a linear regression analysis. (r=-0.076, ns p=0.618, N=45 blood samples).

**Supplementary Figure 10.**

**A.** Gender has on significant influence on Bcl-xS/Bcl-xL expression ratio obtained from urine sediment samples: cDNA was obtained from RNA, extracted from overnight urine sample sediments.. RTPCR was performed using Bcl-x primers to amplify the long and short isoforms. The bioanalyser was used to quantify isoform concentration and Bclxs/Bclxl ratio was calculated. The Bcl-xS/Bcl-xL expression ratio was plotted based on patient’s gender. The data was statistically analysed by a Mann-Whitney test. (mean= 2.52; ns p=0.747; N=15 patients; SEM ±0.501); Female (mean= 2.15; ns p=0.747; N=14 patients; SEM ±0.388).

**B.** There is no significant difference in Bclxs/BclxL expression ratio obtained from leucocytes between males and females: cDNA was obtained from RNA, extracted from leucocytes from blood samples. RTPCR was performed using Bcl-x primers to amplify the long and short isoforms. The bioanalyser was used to quantify isoform concentration and Bclxs/Bclxl ratio was calculated. The Bcl-xS/Bcl-xL expression ratio was plotted based on patient’s gender. The data was statistically analysed by a Mann-Whitney test. Male (mean= 2.78; ns p=0.453; N=30 patients; SEM ±0.512); Female (mean= 2.32; ns p=0.453; N=15 pateints; SEM ±0.592)

**Supplementary Figure 11.**

**Ai. IL-6 increases the pro-apoptotic Bcl-xS band intensity in a concentration-dependent manner.** HEK293 cells were treated with or without recombinant human IL-6 at concentrations 0 ng, 0.1 ng, 1 ng, 10 ng, 25 ng or 50 ng and RNA was extracted after 48 hrs and made into cDNA and RT-PCR performed for Bcl-x isoforms.

**Aii. IL-6 increases Bcl-xs/Bcl-xL ratio.** Bands from **Ai.** were analyzed by densitometry using image j.

**B. IL-6 increases apoptosis in a dose-wise manner.** HEK293 cells were treated with or without recombinant human IL-6 at concentrations 0 ng, 0.1 ng, 1 ng, 10 ng, 25 ng or 50 ng and a JC-10 assay was performed after 24 hrs of treatment. The grapth represents a ratio of apoptotic cells (green) to normal cells (red). 0.1 ng/ml (mean=1.75, ±0.27 SEM, nsp=0.092, N=17 repeats), 1 ng/ml (mean=2.03, ±0.37 SEM, *p=0.028, N=18 repeats), 10 ng/ml (mean=2.38, ±0.37 SEM, ***p=0.0009, N=16 repeats), 25 ng/ml (mean=2.23, ±0.42 SEM, **p=0.007, N=15 repeats) and 50 ng/ml (mean=3.23, ±0.53 SEM, ****p<0.0001, N=18 repeats). Data were statistically analysed using the Kruskal-Wallis test and a multiple comparison post-hoc analysis

**Supplementary Figure 12. SF3B1 expression is not regulated by IL-6 in GEnCs.** GEnCs were treated for 6 h (A) and 24 h (B) with 0, 1, or 25 ng/ml IL-6. There was no significant change in the mRNA expression of SF3B1 after either 6 or 24 h in response to IL-6 treatment (n=3; p=ns as assessed by Kruskal-Wallis).

