## Supplementary material for "The apoptosis gene *BCL-X* splice isoforms have opposing effects in diabetic kidney disease: potential treatment target and prognostic value": Table 1

**Table 1.** MISO analysis of interesting alternative splicing events significantly dysregulated in GEnCs exposed to diabetic conditions (glucose soup) in comparison to a normal glucose and osmotic control.*

| **Gene** | **Event** | **Description** | **Protein** | **Validated with RT-PCR** |
| --- | --- | --- | --- | --- |
| DOK1 | MXE | Scaffold protein. Negative regulator of insulin signalling pathway | Truncated C-terminus missing essential tyrosine docking sites | P<0.05 |
| IRAK1 | MXE | IL-1 receptor associated kinase – we see an increase in the 1b isoform | Lacks an internal segment – IRAK1b does not undergo modification or autophosphorylation and is resistant to degradation | P<0.05 |
| MDM2 | MXE | Proto-oncogene – nuclear localised E3 ubiquitin ligase. | Isoform lacks nuclear localisation and export sequences | P<0.05 |
| TRA2A | MXE | Regulates pre-mRNA splicing | Unknown effect | - |
| ARAP1 | IR | Rho GTPase and GPCR signalling – kidney hypertrophy and apoptosis | Unknown effect | P<0.05 |
| GSTK1 | IR | Enzyme required for cellular detoxification | Shorter isoform – lacks internal segment | - |
| MCRIP1 | IR | MAP kinase signalling pathway involved in epithelial-mesenchymal transition | Shorter isoform may alter ERK signalling | P<0.05 |
| BCL-X | A5SS | Apoptosis | Increase in the apoptotic Bcl-xS | P<0.05 |

*excluding non-protein coding genes. SE, skipped exon; MXE, mutually exclusive exons; A3SS, alternative 3’ splice site; A5SS, alternative 5’ splice site; IR, intron retention
