## Supplementary material for "The apoptosis gene *BCL-X* splice isoforms have opposing effects in diabetic kidney disease: potential treatment target and prognostic value": Table 2

**Table 2.** Patient Characteristics

| Characteristics |  |  |  |
| --- | --- | --- | --- |
|  | **Control** | **Normoalbuminuric** | **Microalbuminuric** |
| N | 27 | 36 | 18 |
| Age years | 70.1 ± 7.3 | 68.4 ± 6.5 | 73.6 ± 5.2 |
| Gender |  |  |  |
| Male | 16(19.8) | 20(24.7) | 15(18.5) |
| Female | 11(13.6) | 16(19.6) | 3(3,7) |
| BMI | 29.7 ± 5.4 | 29.3 ± 4.4 | 27.0 ± 0.4 |
| HbA1c | 41.5 ± 7.3 | 58.0 ± 15.5 | 62.2 ± 10.2 |
| Blood glucose | 5.6 ± 1.3 | 8.1 ± 2.6 | 8.6 ± 1.9 |
| SBP | 131.6 ± 9.6 | 133.5 ± 14.5 | 136.4 ± 29.1 |
| DBP | 80.9 ± 11.77 | 82.6 ± 14.2 | 77.8 ± 11.8 |

Abbreviations: N, total number of patients; n, sample number; BMI, body mass index; SBP, systolic blood pressure; DBP, diastolic blood pressure; HbA1c, glycated haemoglobin. Data are presented as ***n (%)*** or ***mean (±SD).*** Percentages were calculated based on the total number of patients.
