## Supplementary data for "The apoptosis gene *BCL-X* splice isoforms have opposing effects in diabetic kidney disease: potential treatment target and prognostic value"

### Slide 1
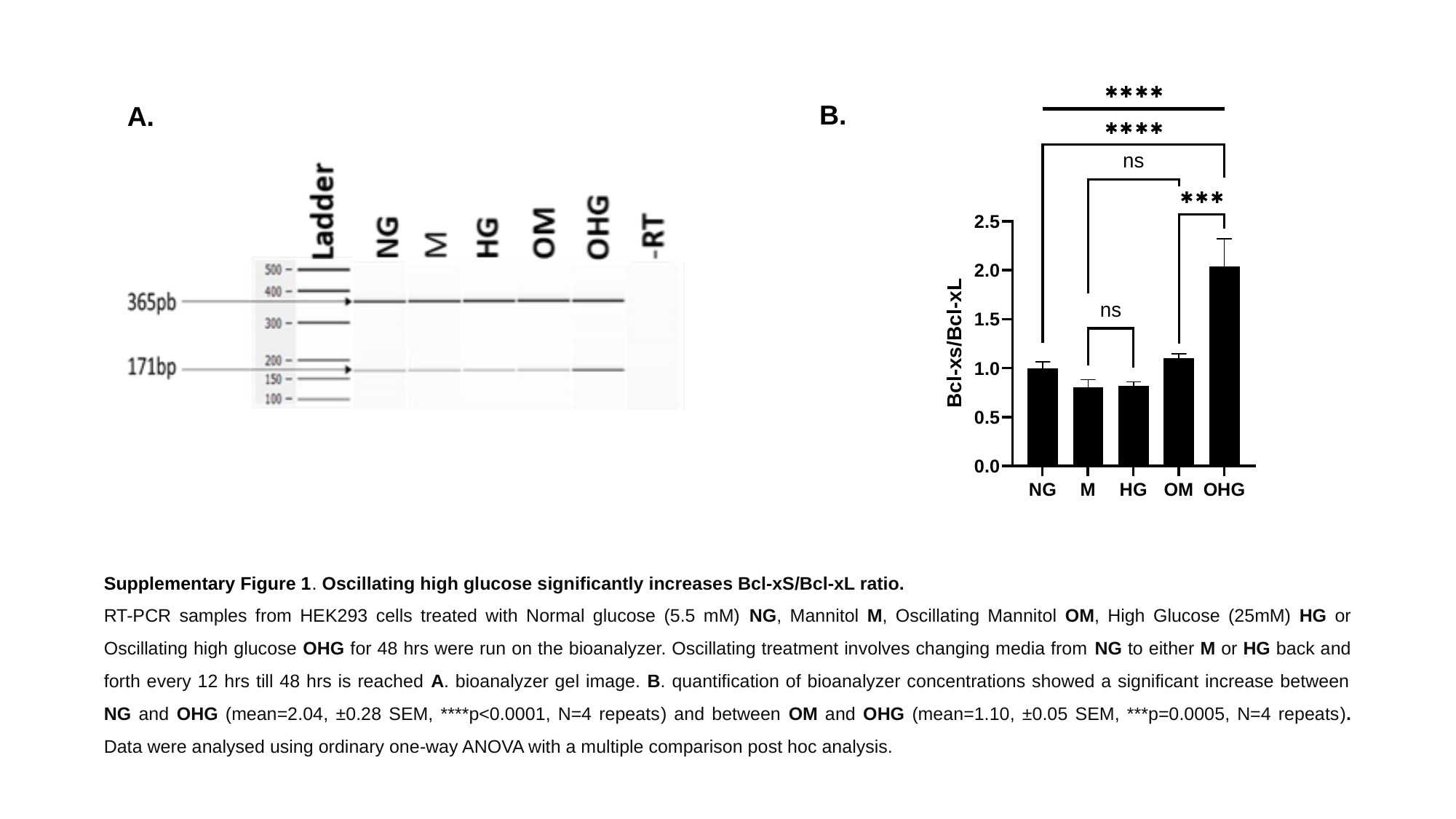

B.
A.
Supplementary Figure 1. Oscillating high glucose significantly increases Bcl-xS/Bcl-xL ratio.
RT-PCR samples from HEK293 cells treated with Normal glucose (5.5 mM) NG, Mannitol M, Oscillating Mannitol OM, High Glucose (25mM) HG or Oscillating high glucose OHG for 48 hrs were run on the bioanalyzer. Oscillating treatment involves changing media from NG to either M or HG back and forth every 12 hrs till 48 hrs is reached A. bioanalyzer gel image. B. quantification of bioanalyzer concentrations showed a significant increase between NG and OHG (mean=2.04, ±0.28 SEM, ****p<0.0001, N=4 repeats) and between OM and OHG (mean=1.10, ±0.05 SEM, ***p=0.0005, N=4 repeats). Data were analysed using ordinary one-way ANOVA with a multiple comparison post hoc analysis.

### Slide 2
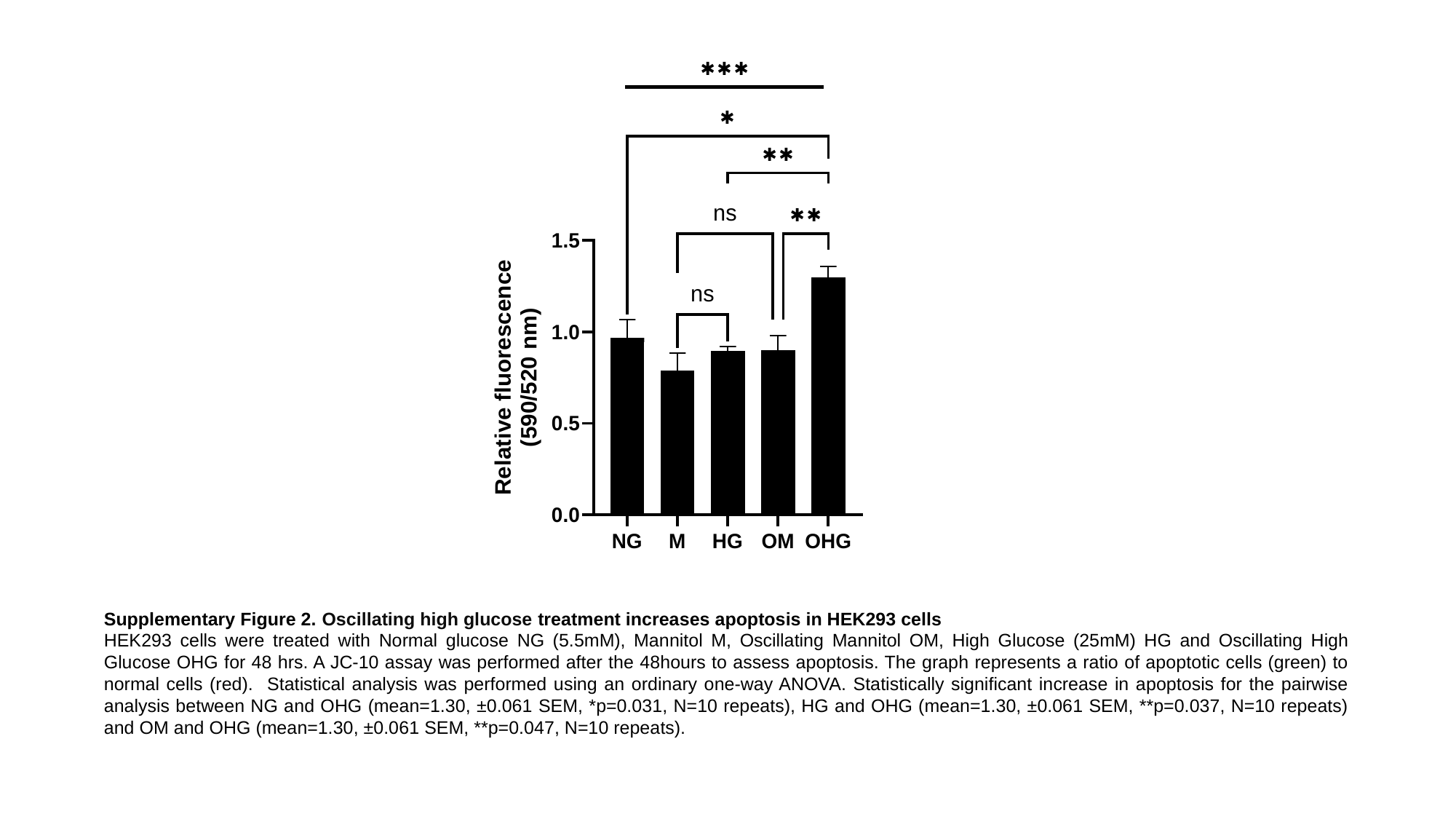

Supplementary Figure 2. Oscillating high glucose treatment increases apoptosis in HEK293 cells
HEK293 cells were treated with Normal glucose NG (5.5mM), Mannitol M, Oscillating Mannitol OM, High Glucose (25mM) HG and Oscillating High Glucose OHG for 48 hrs. A JC-10 assay was performed after the 48hours to assess apoptosis. The graph represents a ratio of apoptotic cells (green) to normal cells (red). Statistical analysis was performed using an ordinary one-way ANOVA. Statistically significant increase in apoptosis for the pairwise analysis between NG and OHG (mean=1.30, ±0.061 SEM, *p=0.031, N=10 repeats), HG and OHG (mean=1.30, ±0.061 SEM, **p=0.037, N=10 repeats) and OM and OHG (mean=1.30, ±0.061 SEM, **p=0.047, N=10 repeats).

### Slide 3
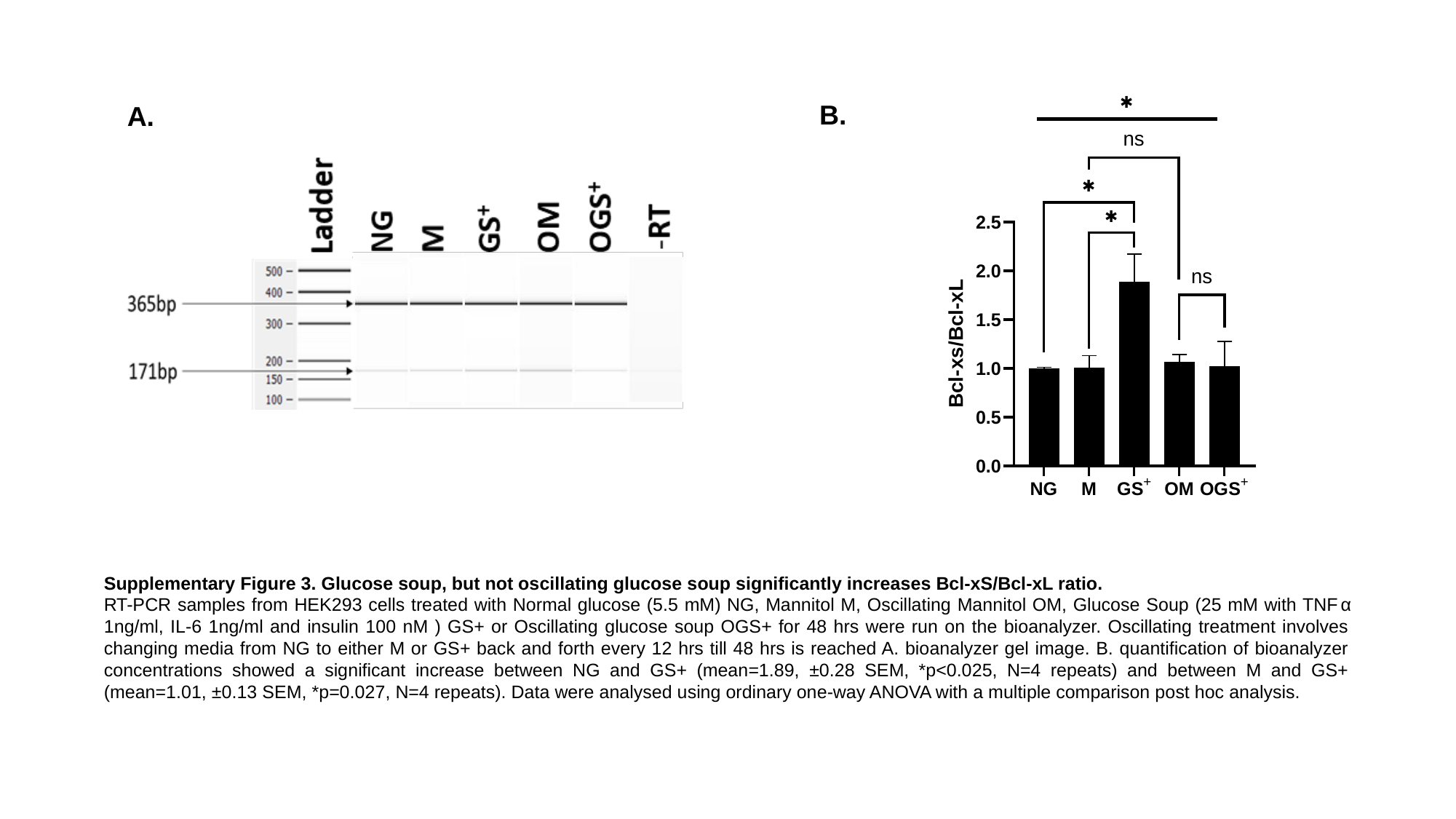

B.
A.
Supplementary Figure 3. Glucose soup, but not oscillating glucose soup significantly increases Bcl-xS/Bcl-xL ratio.
RT-PCR samples from HEK293 cells treated with Normal glucose (5.5 mM) NG, Mannitol M, Oscillating Mannitol OM, Glucose Soup (25 mM with TNFα 1ng/ml, IL-6 1ng/ml and insulin 100 nM ) GS+ or Oscillating glucose soup OGS+ for 48 hrs were run on the bioanalyzer. Oscillating treatment involves changing media from NG to either M or GS+ back and forth every 12 hrs till 48 hrs is reached A. bioanalyzer gel image. B. quantification of bioanalyzer concentrations showed a significant increase between NG and GS+ (mean=1.89, ±0.28 SEM, *p<0.025, N=4 repeats) and between M and GS+ (mean=1.01, ±0.13 SEM, *p=0.027, N=4 repeats). Data were analysed using ordinary one-way ANOVA with a multiple comparison post hoc analysis.

### Slide 4
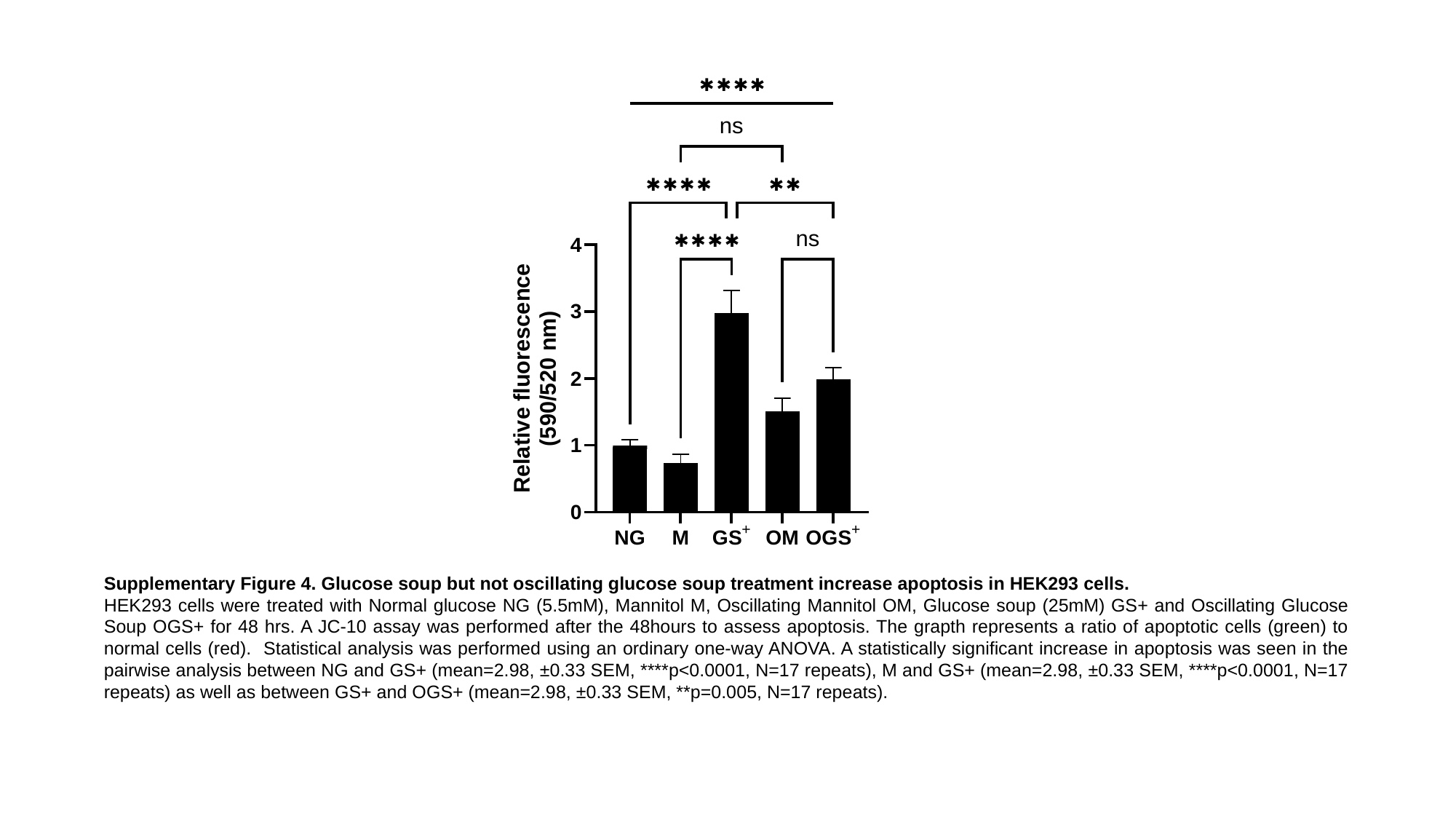

Supplementary Figure 4. Glucose soup but not oscillating glucose soup treatment increase apoptosis in HEK293 cells.
HEK293 cells were treated with Normal glucose NG (5.5mM), Mannitol M, Oscillating Mannitol OM, Glucose soup (25mM) GS+ and Oscillating Glucose Soup OGS+ for 48 hrs. A JC-10 assay was performed after the 48hours to assess apoptosis. The grapth represents a ratio of apoptotic cells (green) to normal cells (red). Statistical analysis was performed using an ordinary one-way ANOVA. A statistically significant increase in apoptosis was seen in the pairwise analysis between NG and GS+ (mean=2.98, ±0.33 SEM, ****p<0.0001, N=17 repeats), M and GS+ (mean=2.98, ±0.33 SEM, ****p<0.0001, N=17 repeats) as well as between GS+ and OGS+ (mean=2.98, ±0.33 SEM, **p=0.005, N=17 repeats).

### Slide 5
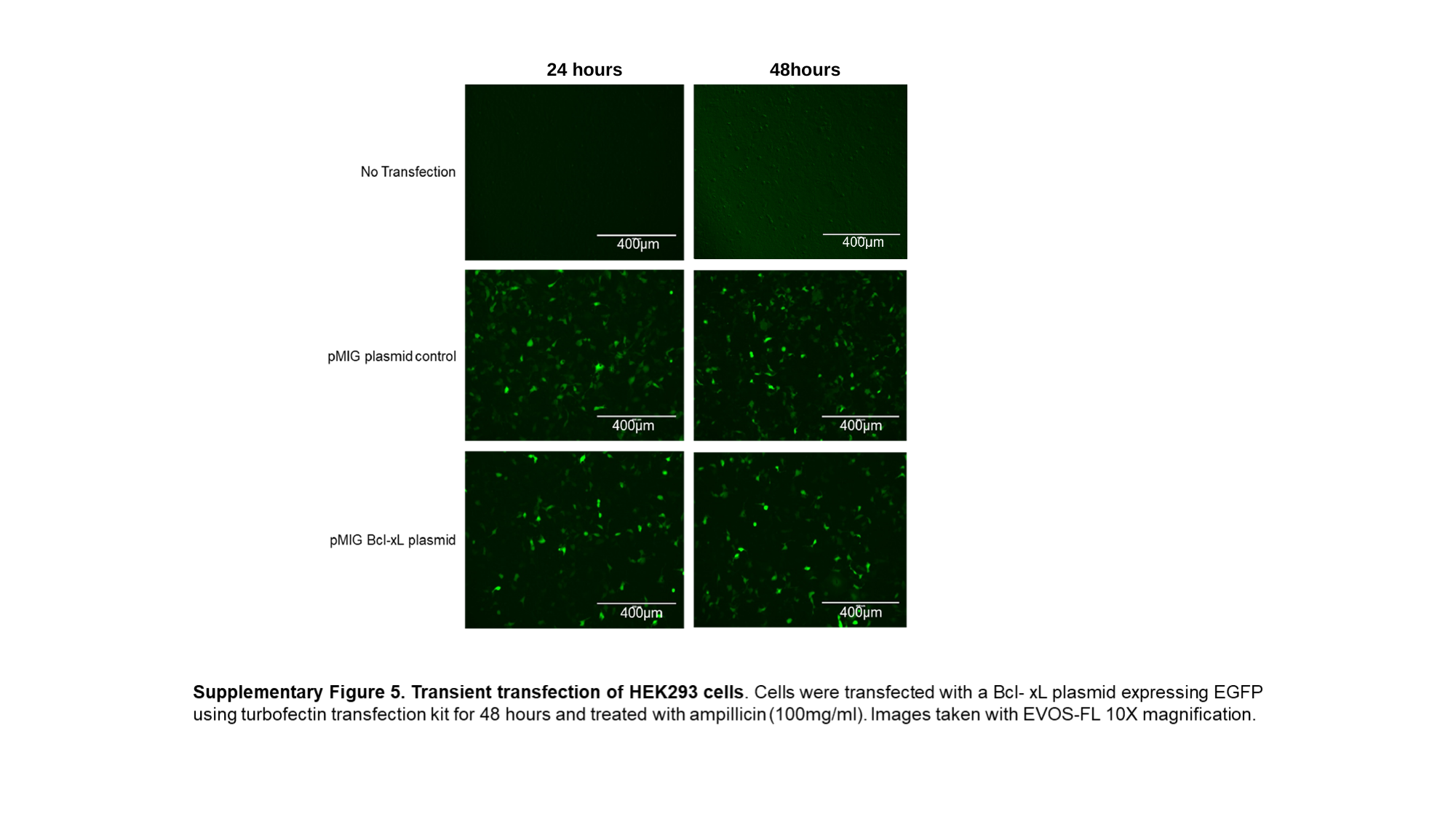

24 hours 48hours

### Slide 6
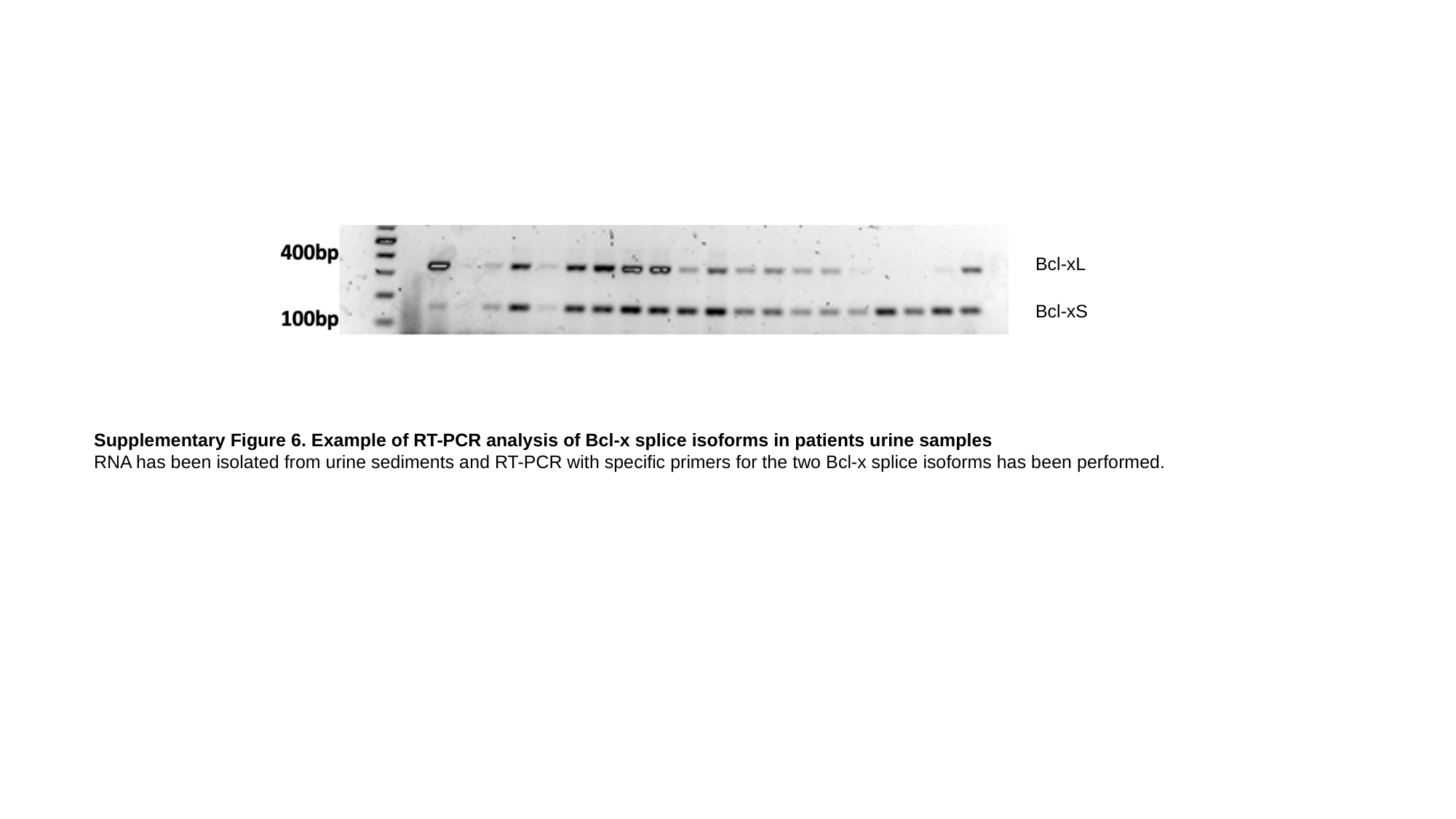

Bcl-xL
Bcl-xS
Supplementary Figure 6. Example of RT-PCR analysis of Bcl-x splice isoforms in patients urine samples
RNA has been isolated from urine sediments and RT-PCR with specific primers for the two Bcl-x splice isoforms has been performed.

### Slide 7
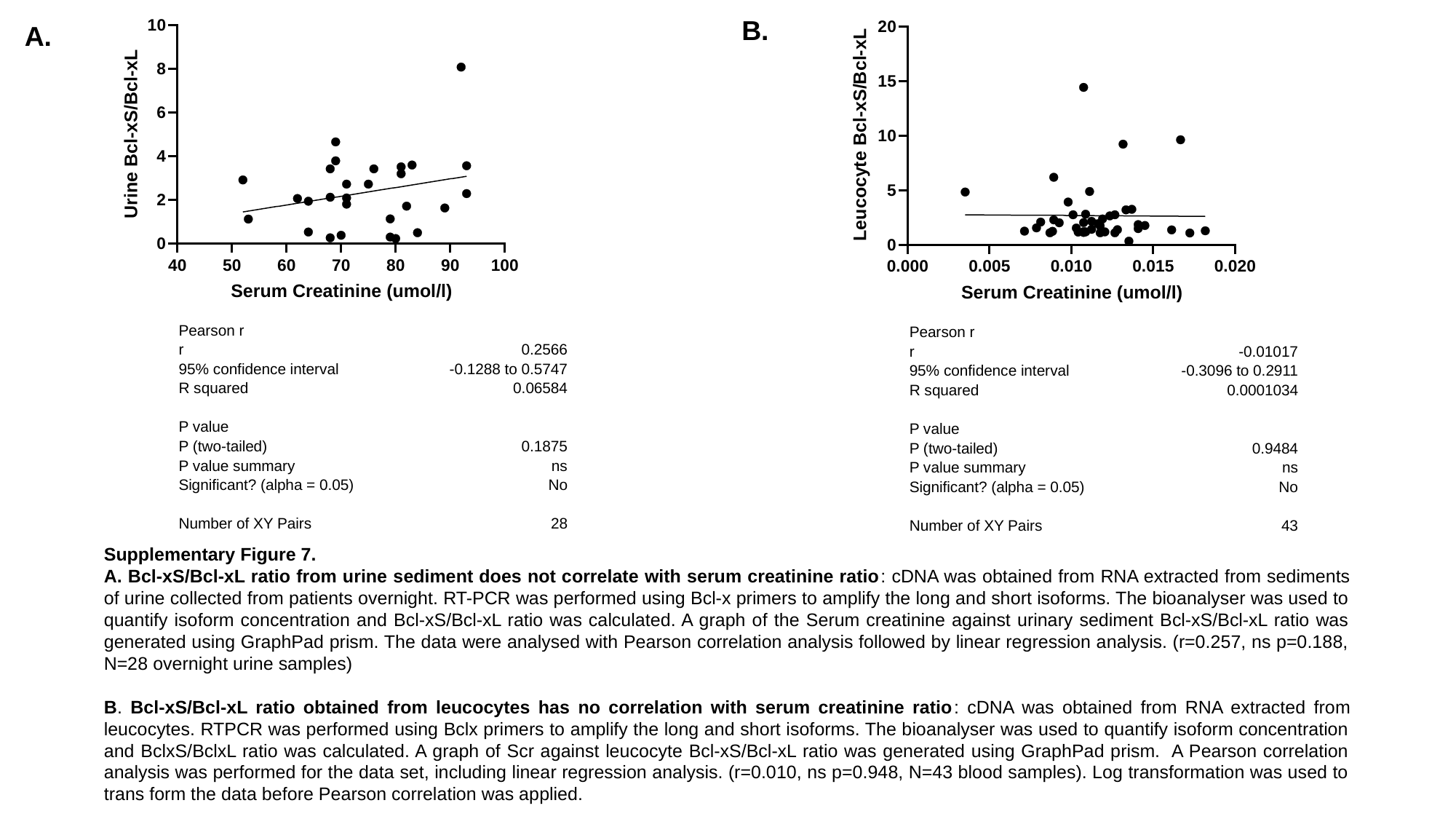

B.
A.
| Pearson r | |
| --- | --- |
| r | 0.2566 |
| 95% confidence interval | -0.1288 to 0.5747 |
| R squared | 0.06584 |
| P value | |
| P (two-tailed) | 0.1875 |
| P value summary | ns |
| Significant? (alpha = 0.05) | No |
| Number of XY Pairs | 28 |
| Pearson r | |
| --- | --- |
| r | -0.01017 |
| 95% confidence interval | -0.3096 to 0.2911 |
| R squared | 0.0001034 |
| P value | |
| P (two-tailed) | 0.9484 |
| P value summary | ns |
| Significant? (alpha = 0.05) | No |
| Number of XY Pairs | 43 |
Supplementary Figure 7.
A. Bcl-xS/Bcl-xL ratio from urine sediment does not correlate with serum creatinine ratio: cDNA was obtained from RNA extracted from sediments of urine collected from patients overnight. RT-PCR was performed using Bcl-x primers to amplify the long and short isoforms. The bioanalyser was used to quantify isoform concentration and Bcl-xS/Bcl-xL ratio was calculated. A graph of the Serum creatinine against urinary sediment Bcl-xS/Bcl-xL ratio was generated using GraphPad prism. The data were analysed with Pearson correlation analysis followed by linear regression analysis. (r=0.257, ns p=0.188, N=28 overnight urine samples)
B. Bcl-xS/Bcl-xL ratio obtained from leucocytes has no correlation with serum creatinine ratio: cDNA was obtained from RNA extracted from leucocytes. RTPCR was performed using Bclx primers to amplify the long and short isoforms. The bioanalyser was used to quantify isoform concentration and BclxS/BclxL ratio was calculated. A graph of Scr against leucocyte Bcl-xS/Bcl-xL ratio was generated using GraphPad prism. A Pearson correlation analysis was performed for the data set, including linear regression analysis. (r=0.010, ns p=0.948, N=43 blood samples). Log transformation was used to trans form the data before Pearson correlation was applied.

### Slide 8
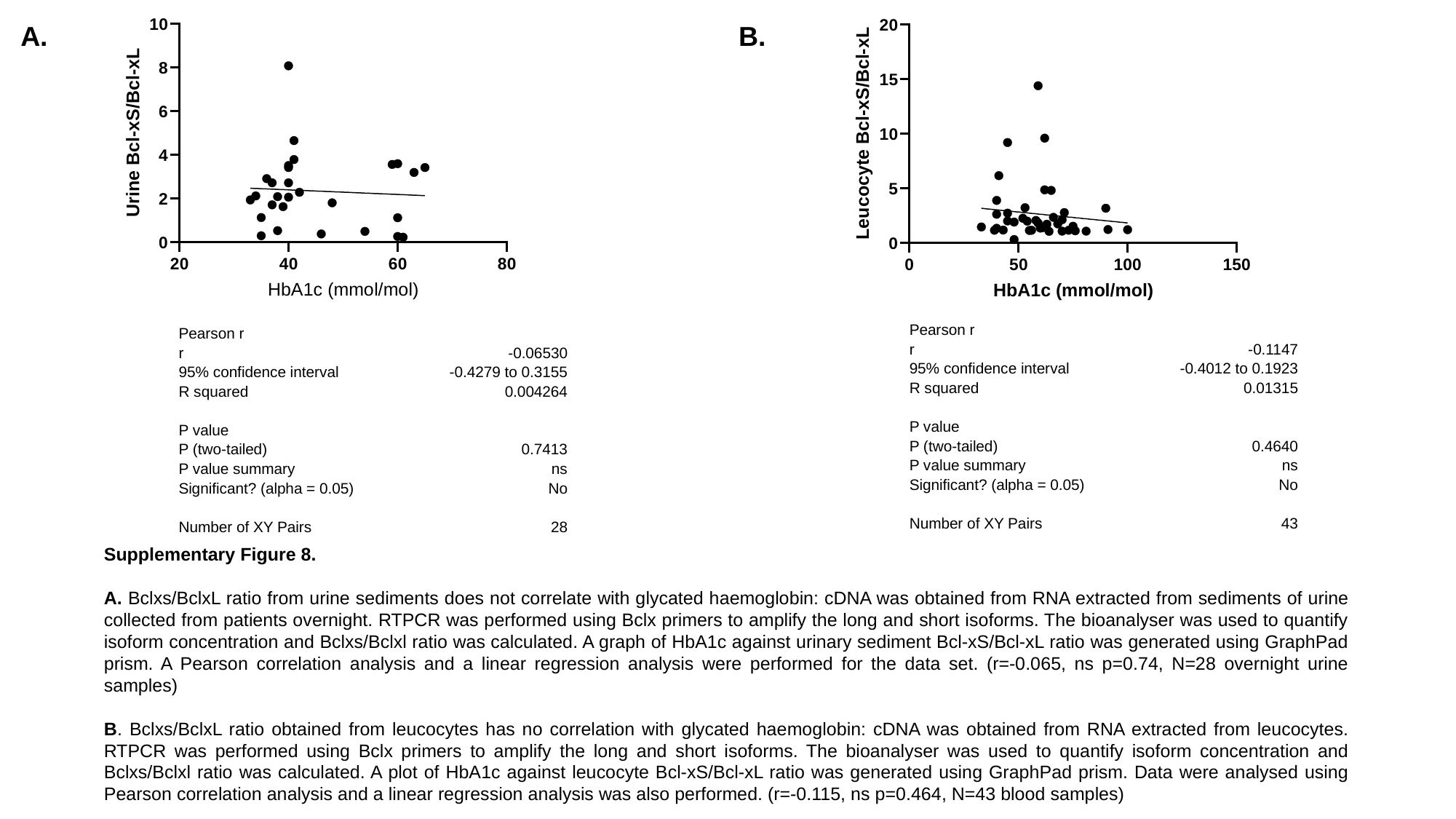

A.
B.
| Pearson r | |
| --- | --- |
| r | -0.1147 |
| 95% confidence interval | -0.4012 to 0.1923 |
| R squared | 0.01315 |
| P value | |
| P (two-tailed) | 0.4640 |
| P value summary | ns |
| Significant? (alpha = 0.05) | No |
| Number of XY Pairs | 43 |
| Pearson r | |
| --- | --- |
| r | -0.06530 |
| 95% confidence interval | -0.4279 to 0.3155 |
| R squared | 0.004264 |
| P value | |
| P (two-tailed) | 0.7413 |
| P value summary | ns |
| Significant? (alpha = 0.05) | No |
| Number of XY Pairs | 28 |
Supplementary Figure 8.
A. Bclxs/BclxL ratio from urine sediments does not correlate with glycated haemoglobin: cDNA was obtained from RNA extracted from sediments of urine collected from patients overnight. RTPCR was performed using Bclx primers to amplify the long and short isoforms. The bioanalyser was used to quantify isoform concentration and Bclxs/Bclxl ratio was calculated. A graph of HbA1c against urinary sediment Bcl-xS/Bcl-xL ratio was generated using GraphPad prism. A Pearson correlation analysis and a linear regression analysis were performed for the data set. (r=-0.065, ns p=0.74, N=28 overnight urine samples)
B. Bclxs/BclxL ratio obtained from leucocytes has no correlation with glycated haemoglobin: cDNA was obtained from RNA extracted from leucocytes. RTPCR was performed using Bclx primers to amplify the long and short isoforms. The bioanalyser was used to quantify isoform concentration and Bclxs/Bclxl ratio was calculated. A plot of HbA1c against leucocyte Bcl-xS/Bcl-xL ratio was generated using GraphPad prism. Data were analysed using Pearson correlation analysis and a linear regression analysis was also performed. (r=-0.115, ns p=0.464, N=43 blood samples)

### Slide 9
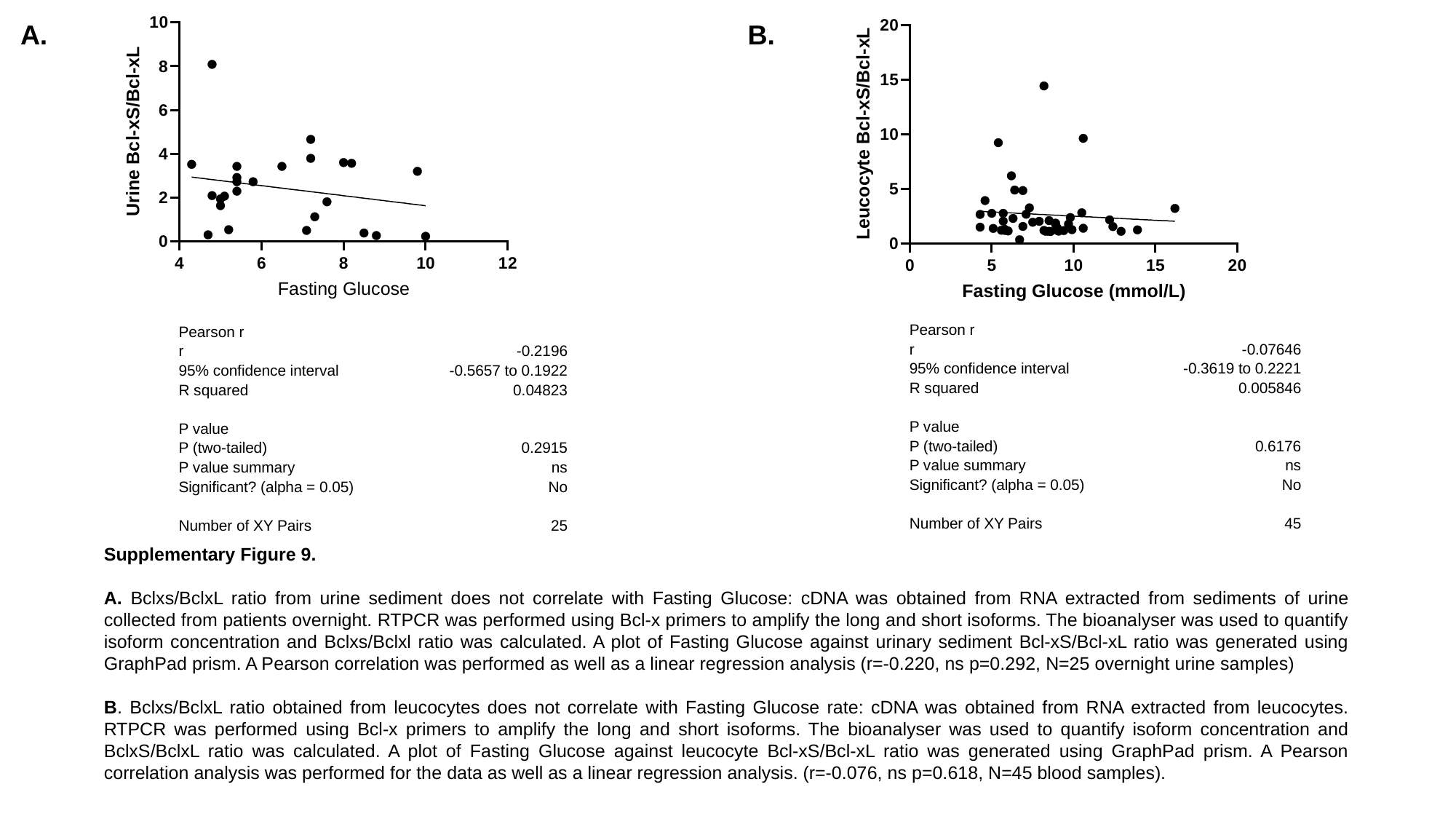

A.
B.
| Pearson r | |
| --- | --- |
| r | -0.07646 |
| 95% confidence interval | -0.3619 to 0.2221 |
| R squared | 0.005846 |
| P value | |
| P (two-tailed) | 0.6176 |
| P value summary | ns |
| Significant? (alpha = 0.05) | No |
| Number of XY Pairs | 45 |
| Pearson r | |
| --- | --- |
| r | -0.2196 |
| 95% confidence interval | -0.5657 to 0.1922 |
| R squared | 0.04823 |
| P value | |
| P (two-tailed) | 0.2915 |
| P value summary | ns |
| Significant? (alpha = 0.05) | No |
| Number of XY Pairs | 25 |
Supplementary Figure 9.
A. Bclxs/BclxL ratio from urine sediment does not correlate with Fasting Glucose: cDNA was obtained from RNA extracted from sediments of urine collected from patients overnight. RTPCR was performed using Bcl-x primers to amplify the long and short isoforms. The bioanalyser was used to quantify isoform concentration and Bclxs/Bclxl ratio was calculated. A plot of Fasting Glucose against urinary sediment Bcl-xS/Bcl-xL ratio was generated using GraphPad prism. A Pearson correlation was performed as well as a linear regression analysis (r=-0.220, ns p=0.292, N=25 overnight urine samples)
B. Bclxs/BclxL ratio obtained from leucocytes does not correlate with Fasting Glucose rate: cDNA was obtained from RNA extracted from leucocytes. RTPCR was performed using Bcl-x primers to amplify the long and short isoforms. The bioanalyser was used to quantify isoform concentration and BclxS/BclxL ratio was calculated. A plot of Fasting Glucose against leucocyte Bcl-xS/Bcl-xL ratio was generated using GraphPad prism. A Pearson correlation analysis was performed for the data as well as a linear regression analysis. (r=-0.076, ns p=0.618, N=45 blood samples).

### Slide 10
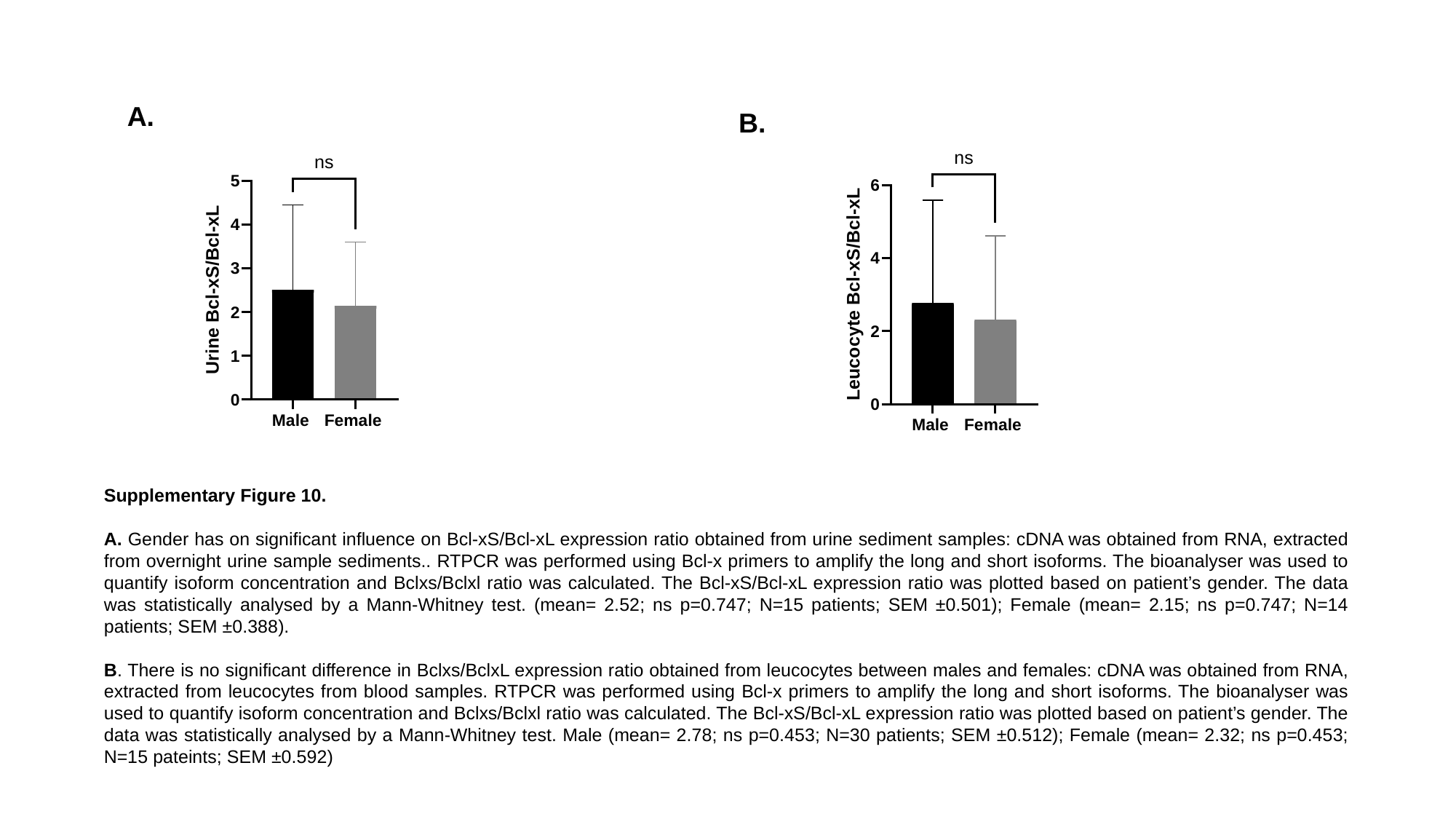

A.
B.
Supplementary Figure 10.
A. Gender has on significant influence on Bcl-xS/Bcl-xL expression ratio obtained from urine sediment samples: cDNA was obtained from RNA, extracted from overnight urine sample sediments.. RTPCR was performed using Bcl-x primers to amplify the long and short isoforms. The bioanalyser was used to quantify isoform concentration and Bclxs/Bclxl ratio was calculated. The Bcl-xS/Bcl-xL expression ratio was plotted based on patient’s gender. The data was statistically analysed by a Mann-Whitney test. (mean= 2.52; ns p=0.747; N=15 patients; SEM ±0.501); Female (mean= 2.15; ns p=0.747; N=14 patients; SEM ±0.388).
B. There is no significant difference in Bclxs/BclxL expression ratio obtained from leucocytes between males and females: cDNA was obtained from RNA, extracted from leucocytes from blood samples. RTPCR was performed using Bcl-x primers to amplify the long and short isoforms. The bioanalyser was used to quantify isoform concentration and Bclxs/Bclxl ratio was calculated. The Bcl-xS/Bcl-xL expression ratio was plotted based on patient’s gender. The data was statistically analysed by a Mann-Whitney test. Male (mean= 2.78; ns p=0.453; N=30 patients; SEM ±0.512); Female (mean= 2.32; ns p=0.453; N=15 pateints; SEM ±0.592)

### Slide 11
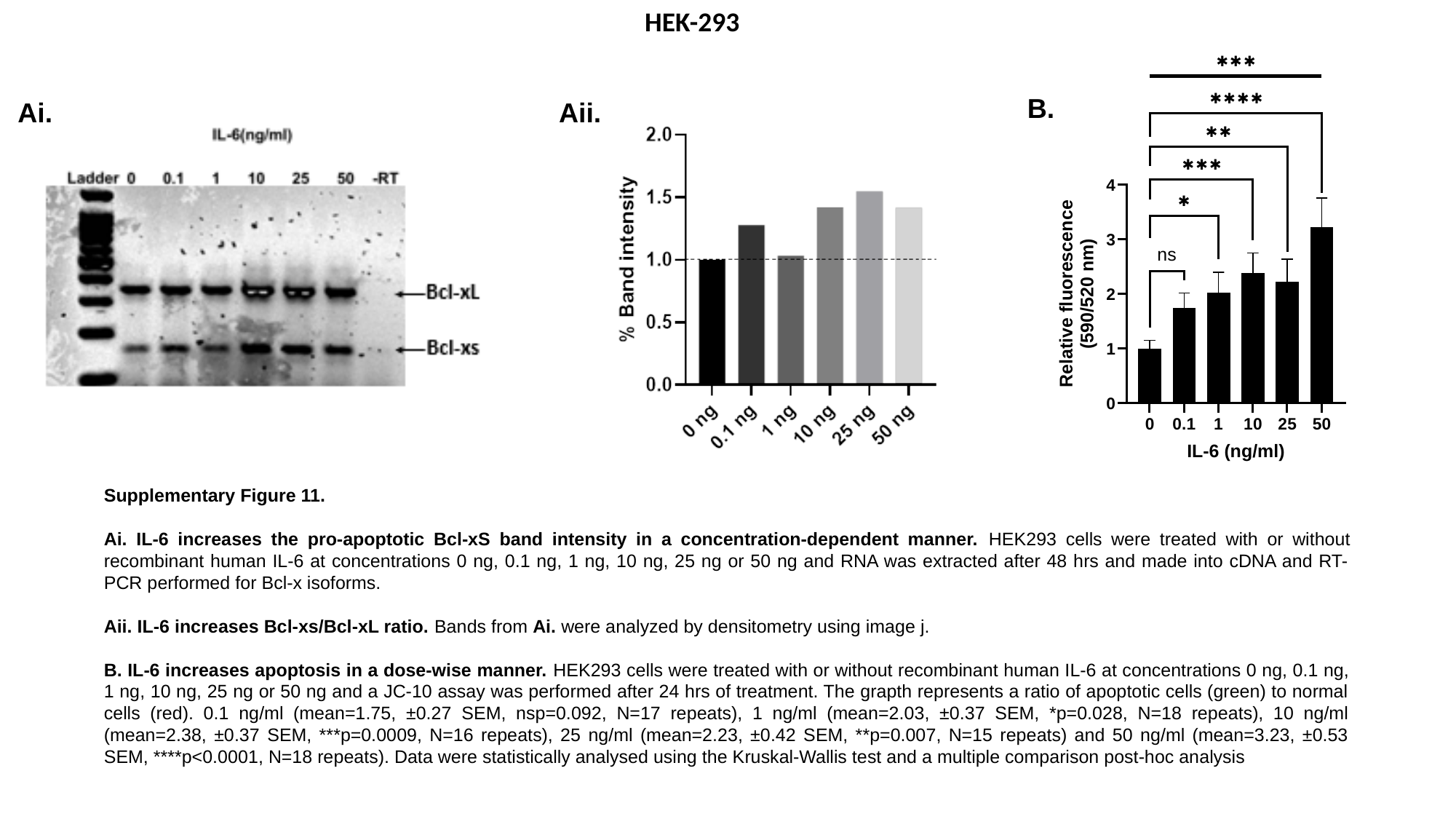

HEK-293
B.
Ai.
Aii.
Supplementary Figure 11.
Ai. IL-6 increases the pro-apoptotic Bcl-xS band intensity in a concentration-dependent manner. HEK293 cells were treated with or without recombinant human IL-6 at concentrations 0 ng, 0.1 ng, 1 ng, 10 ng, 25 ng or 50 ng and RNA was extracted after 48 hrs and made into cDNA and RT-PCR performed for Bcl-x isoforms.
Aii. IL-6 increases Bcl-xs/Bcl-xL ratio. Bands from Ai. were analyzed by densitometry using image j.
B. IL-6 increases apoptosis in a dose-wise manner. HEK293 cells were treated with or without recombinant human IL-6 at concentrations 0 ng, 0.1 ng, 1 ng, 10 ng, 25 ng or 50 ng and a JC-10 assay was performed after 24 hrs of treatment. The grapth represents a ratio of apoptotic cells (green) to normal cells (red). 0.1 ng/ml (mean=1.75, ±0.27 SEM, nsp=0.092, N=17 repeats), 1 ng/ml (mean=2.03, ±0.37 SEM, *p=0.028, N=18 repeats), 10 ng/ml (mean=2.38, ±0.37 SEM, ***p=0.0009, N=16 repeats), 25 ng/ml (mean=2.23, ±0.42 SEM, **p=0.007, N=15 repeats) and 50 ng/ml (mean=3.23, ±0.53 SEM, ****p<0.0001, N=18 repeats). Data were statistically analysed using the Kruskal-Wallis test and a multiple comparison post-hoc analysis

### Slide 12
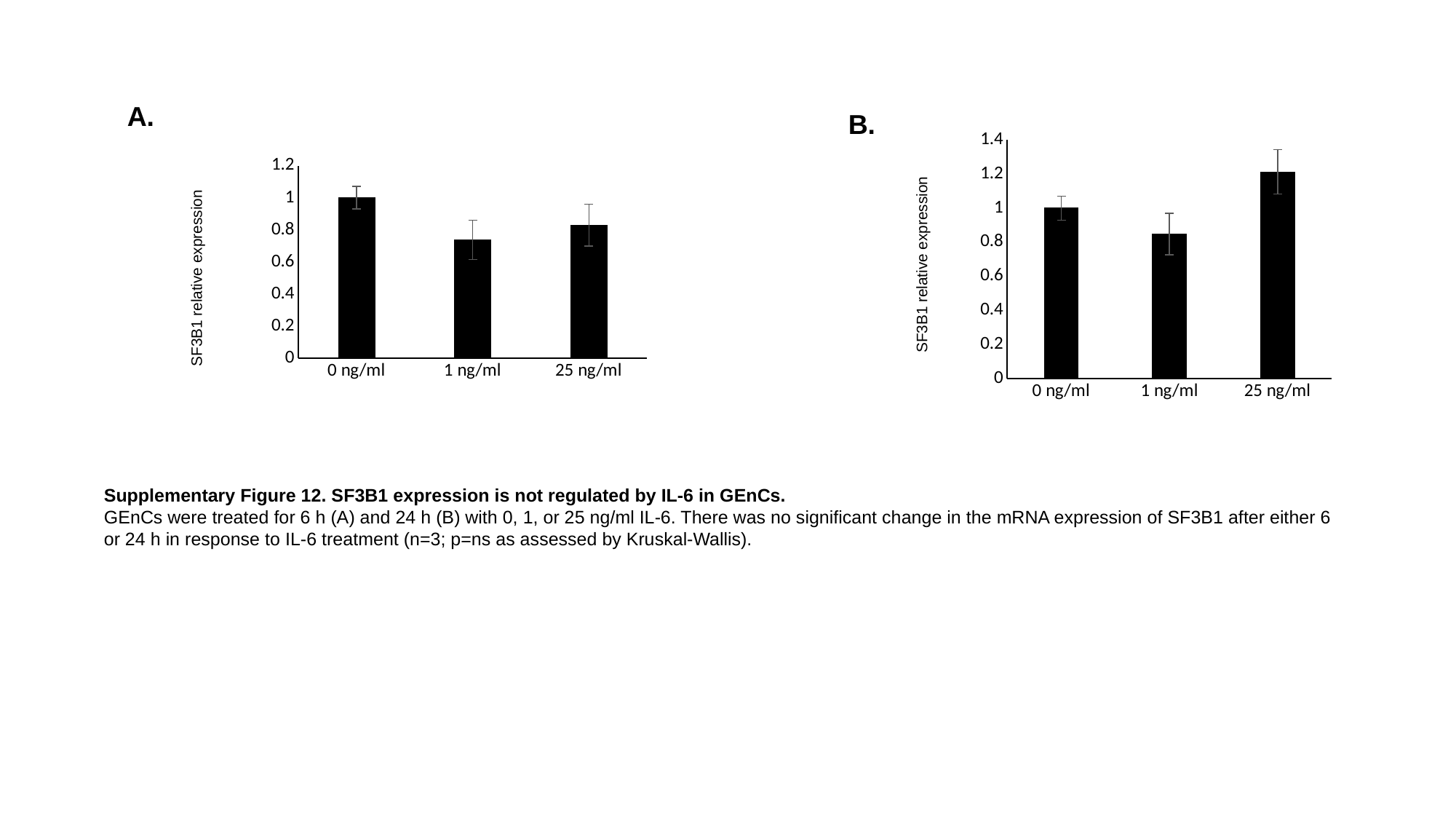

A.
B.
#### Chart
| Category | |
|---|---|
| 0 ng/ml | 1.0 |
| 1 ng/ml | 0.8469635992601473 |
| 25 ng/ml | 1.2120865814277724 |
#### Chart
| Category | |
|---|---|
| 0 ng/ml | 1.0 |
| 1 ng/ml | 0.7379271904853937 |
| 25 ng/ml | 0.8302104081219511 |SF3B1 relative expression
SF3B1 relative expression
Supplementary Figure 12. SF3B1 expression is not regulated by IL-6 in GEnCs.
GEnCs were treated for 6 h (A) and 24 h (B) with 0, 1, or 25 ng/ml IL-6. There was no significant change in the mRNA expression of SF3B1 after either 6 or 24 h in response to IL-6 treatment (n=3; p=ns as assessed by Kruskal-Wallis).
